# Loss of functional heterogeneity along the CA3 transverse axis in aging

**DOI:** 10.1101/2021.09.14.460329

**Authors:** Heekyung Lee, Zitong Wang, Arjuna Tillekeratne, Nick Lukish, Scott Zeger, Michela Gallagher, James J. Knierim

## Abstract

Age-related deficits in pattern separation have been postulated to bias the output of hippocampal memory processing toward pattern completion, which can cause deficits in accurate memory retrieval. While the CA3 region of the hippocampus is often conceptualized as a homogeneous network involved in pattern completion, growing evidence demonstrates a functional gradient in CA3 along the transverse axis, as pattern-separated outputs (dominant in the more proximal CA3) transition to pattern-completed outputs (dominant in the more distal CA3). We examined the neural representations along the CA3 transverse axis in young (Y), aged memory-unimpaired (AU), and aged memory-impaired (AI) rats when different changes were made to the environment. Functional heterogeneity in CA3 was observed in Y and AU rats when the environmental similarity was high (altered cues or altered environment shapes in the same room), with more orthogonalized representations in proximal CA3 than in distal CA3. In contrast, AI rats showed reduced orthogonalization in proximal CA3 but showed normal (i.e., generalized) representations in distal CA3, with little evidence of a functional gradient. Under experimental conditions when the environmental similarity was low (different rooms), representations in proximal and distal CA3 remapped in all rats, showing that CA3 of AI rats is able to encode distinctive representations for inputs with greater dissimilarity. These experiments support the hypotheses that the aged-related bias towards hippocampal pattern completion is due to the loss in AI rats of the normal transition from pattern separation to pattern completion along the CA3 transverse axis.

## Introduction

Aging increases an organism’s susceptibility to memory interference (Ebert and Anderson, 2009; Emery et al., 2008; Lustig and Hasher, 2001), which occurs when similar memories compete with each other and impair the correct retrieval of any one of those memories. To minimize memory interference and maximize storage capacity in a distributed memory system, computational models hypothesized that the hippocampus performs two complementary processes: pattern separation (the ability to orthogonalize similar input patterns to create less overlapping output patterns) and pattern completion (the ability to retrieve stored output patterns when presented with partial or degraded input patterns) (Marr, 1971; McClelland and Goddard, 1996; McNaughton and Morris, 1987; O’Reilly and McClelland, 1994; Rolls and Treves, 1998; Treves and Rolls, 1992). Most formulations of these models propose that the dentate gyrus (DG) performs pattern separation and the CA3 region performs pattern completion (although other models of DG-CA3 circuit function have also been proposed; e.g., (Hainmueller and Bartos, 2020; Lee and Jung, 2017; Lisman, 1999)).

Place cell remapping is often used as a proxy to study pattern separation. Place cells are hippocampal neurons that fire when an animal occupies a specific location in an environment (O’Keefe and Dostrovsky, 1971) and are proposed to create a cognitive map that stores experiences associated with that environment (O’Keefe and Nadel, 1978). Place cells undergo remapping when changes are made to the environment or when the animal is moved to a different environment (Muller and Kubie, 1987), as well as when the animal changes its behavior or internal state (Ferbinteanu and Shapiro, 2003; Kennedy and Shapiro, 2009; Markus et al., 1995; Moita et al., 2004; Wood et al., 2000). In remapping, place fields can appear, disappear, change firing locations, or change firing rates (Anderson and Jeffery, 2003; Knierim, 2003; Leutgeb et al., 2005; Muller and Kubie, 1987). Thus, remapping creates dissimilar spatial maps for distinct experiences or contexts that can be stored as different memories.

Converging studies from rodents and humans have demonstrated an age-related decline in pattern separation abilities. Rodent studies have shown that CA3 place cells of aged rats failed to encode changes in the environment, demonstrating a rigidity in their representations (Robitsek et al., 2015; Wilson et al., 2005). Similarly, in human fMRI studies, representational rigidity in older adults’ DG/CA3 region was linked to mnemonic discrimination deficits (Lacy et al., 2011; Yassa et al., 2010, 2011). Studies of aged CA3 have indicated that hyperactivity contributes to pattern separation deficits by strengthening the autoassociative CA3 network to favor enhanced pattern completion (Haberman et al., 2017; Lee et al., 2021; Maurer et al., 2017; Robitsek et al., 2015; Thomé et al., 2016; Wilson et al., 2005; Yassa et al., 2011). Accordingly, treatments to reduce hyperactivity with the antiepileptic drug, levetiracetam, improved memory performance in aged rats (Haberman et al., 2017; Koh et al., 2010; Robitsek et al., 2015) and in amnestic mild cognitive impairment patients (Bakker et al., 2012, 2015), suggesting that normalizing CA3 hyperactivity restored the balance between pattern separation and pattern completion.

Anatomical and functional heterogeneity along the CA3 transverse axis suggests that proximal CA3 supports pattern separation and distal CA3 supports pattern completion (Hunsaker et al., 2008; Ishizuka et al., 1990; Lee et al., 2015; Lu et al., 2015; Marrone et al., 2014; Nakamura et al., 2013; Sun et al., 2017; Witter, 2007). This gradient is reinforced by lesion studies in rats. Proximal CA3 lesions, but not intermediate or distal CA3 lesions, led to a deficit in the ability to discriminate small changes in the distance between two objects (Hunsaker et al., 2008). In contrast, lesions to intermediate and distal CA3 showed deficits in pattern completion on a delayed matching-to-place task (Gold and Kesner, 2005). Because age-related hyperactivity appears restricted towards proximal CA3, while potential hypoactivity is present towards distal CA3 (Lee et al., 2021), the age-related bias from pattern separation to pattern completion may be specific to the proximal CA3 region. However, there have been no direct studies examining the neural representations along the CA3 transverse axis in aging. The present study aims to determine how age-related dysfunctions may result from an imbalance of the normal gradient of pattern separation and pattern completion along the CA3 transverse axis.

## Results

### Histology and experimental procedure

Multielectrode recordings were made along the dorsal CA3 transverse axis (Fig. 1A, Supp. Fig.1) in 4 young and 14 aged rats. All rats were behaviorally characterized in the Morris water maze (see Methods), and the aged rats were categorized as learning impaired (AI; n = 7) or learning unimpaired (AU; n = 7) based on the pre-determined learning index (LI) score of 240, which represents 2 standard deviations from the mean for typical Y rats tested over many experiments (Branch et al., 2019; Gallagher et al., 1993) (Fig. 1B). Three experiments were performed. In the *cue-conflict experiment*, all rats were trained to run clockwise (CW) on a circular track located in a curtained room (Fig. 1C). Four different local textures tiled the surface of the circular track and multiple global cues were placed along the curtained room (Knierim, 2002). Three standard (STD) sessions with a fixed local and global cue configuration were interleaved with two cue-mismatch (MIS) sessions, in which the track was rotated in a counterclockwise (CCW) direction and the global cues in a CW direction. In the *two-shape experiment*, subsets of the rats (3 Y, 6 AU, and 6 AI) were trained to forage in circle and square enclosures placed at a constant location in room A. In the *two-room experiment*, the same subsets of rats were taken to room B (after the four sessions of alternating square and circle in room A) and two more sessions were recorded in the circle and square enclosures placed at a constant location in room B.

**Figure 1.**
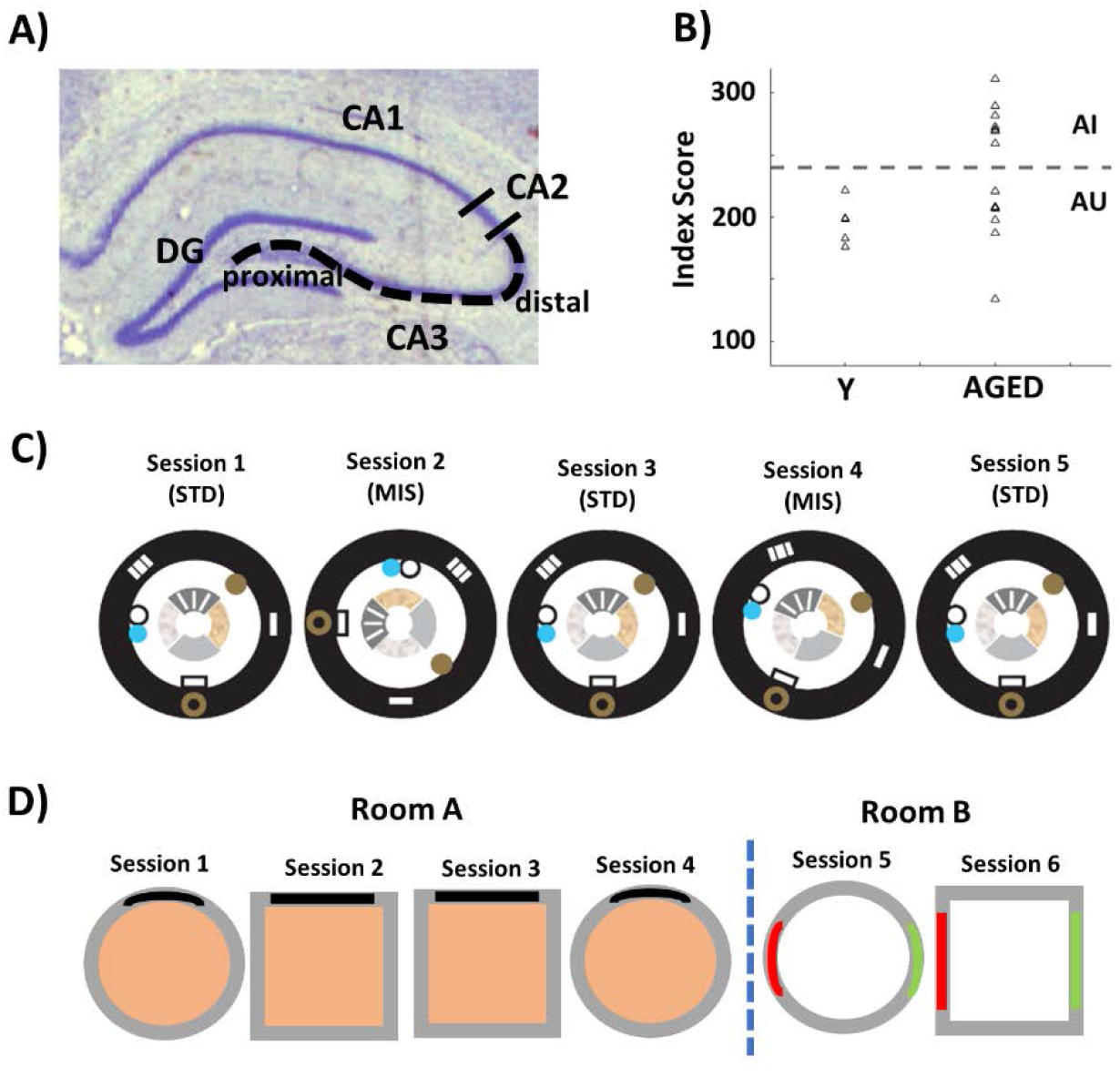
CA3 recording locations and experimental procedures. **(A)** Recordings were made along the CA3 transverse axis. **(B)** A learning index score was derived from measures of proximity to the goal location during probe trials interpolated throughout the water maze training, with lower scores indicating more accurate performance. Aged rats that performed on par with young rats (below the dotted line, representing 2 S.D. above the mean of Y rats based on a large population study) were designated as aged-unimpaired (AU), and those that performed more poorly than young rats (above the dotted line) were designated as aged-impaired (AI), according to established criteria (Gallagher et al., 1993). The gap in LI values between AI and AU rats in the present data is purely the result of sampling error; in the underlying population of aged rats, the values of LI form a smooth transition between AU and AI rats (Gallagher et al., 1993). **(C)** In the local-global cue mismatch experiment, recording sessions consisted of three standard (STD) sessions interleaved with two mismatch (MIS) sessions. In the STD sessions, the local cues on the track (denoted by the inner ring with four different textures) and the global cues along the curtain at the periphery (denoted by the black outer ring) were arranged in a fixed configuration that the rat had experienced during training periods. In the MIS sessions, the global cues were rotated in a clockwise direction and the local cues were rotated in a counterclockwise direction by the same amount, for a net mismatch of 90°, 135°, or 180°. **(D)** In the two-shape experiment, recording sessions consisted of 4 sessions (two sessions in a circle and two sessions in a square) in room A. In the two-room experiment, after 4 sessions in room A, two additional sessions (one session each in circle and square) were recorded in room B. In room A, there was a striped cue card on the north wall of the circle and square enclosures placed above a brown paper floor. In room B, there were two cue cards (a brown cue card on the west and a gray cue card on the east) on the walls of the enclosures placed above a white paper floor.

### Individual cell responses in the cue-conflict manipulations along the CA3 transverse axis

In the cue-conflict experiment, individual cell responses to the cue manipulations were categorized into five groups, similar to previously published studies using the same task (Lee et al., 2015, 2004; Neunuebel and Knierim, 2014) (Fig. 2A). Classifications were based on each cell’s response in the MIS session compared to the cell’s firing properties in the immediately preceding STD session. For simplicity, the 5 categories were condensed into 2 categories, such that if a cell met the place-field inclusion criteria (spatial information score > 0.75, significance p < 0.01, number of on-track spikes > 50) in both STD and MIS sessions, the cell was classified as a “Rotate” cell. If a cell met the place-field inclusion criteria in either the STD or MIS session, but not in both, the cell was classified as a “Remap” cell. Because previous studies of CA3 in this cue-mismatch experiment have shown that the large majority of Rotate cells are controlled coherently by the local set of cues (Knierim, 2002; Lee et al., 2015, 2004; Neunuebel and Knierim, 2014), we consider this class as approximately indicating the degree to which the hippocampus maintains the same representation between the two sessions (i.e., these same cells represent both the STD and MIS conditions, under the control of the local frame of reference). In contrast, cells that are classified as Remap indicate the degree to which the hippocampus creates different representations between the environments (i.e., different populations of cells represent the STD and MIS environments). For our initial analysis of cell counts, we subdivided CA3 into 3 regions, corresponding approximately to classic definitions of CA3a, CA3b, and CA3c (Lorente de No, 1934).

**Figure 2.**
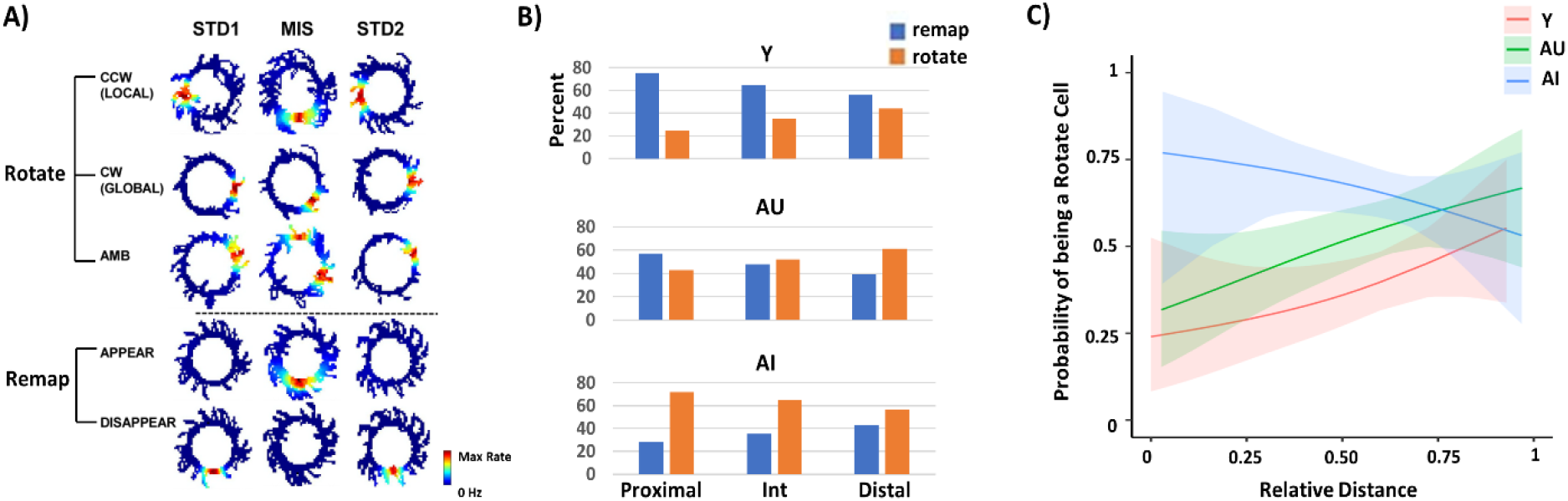
Responses to the cue manipulations differ among the groups along the CA3 transverse axis. **(A)** Examples of five different response types observed in CA3. Cells that showed CCW rotation with the local cues, CW rotation with the global cues, and ambiguous (AMB) rotation were categorized as “Rotate” cells. For example, compared to the location of the place field in STD1, the location of the place field in MIS rotated in either a CCW or a CW direction. In AMB rotation, a single place field in STD1 split into two fields in MIS, with one field showing CCW rotation and the other field showing CW rotation. Cells with place fields that APPEAR in MIS or DISAPPEAR in MIS were categorized as “Remap” cells. **(B)** Classification of responses in different CA3 subregions are different in age groups. As was done in our prior study (Lee et al., 2021), CA3 was subdivided to match approximately the classic definitions of CA3a, CA3b, and CA3c as proximal (40% of the length along the transverse axis), intermediate (the next 30%), and distal (the final 30%). Y and AU rats showed a similar trend of a decreasing proportion of Remap cells from proximal to distal CA3, with an increasing proportion of Rotate cells from proximal to distal CA3. In contrast, AI rats showed an opposite trend, with a decreasing proportion of Rotate cells from proximal to distal CA3 and an increasing proportion of Remap cells from proximal to distal CA3. **(C)** Logistic regression model showing the probability that cells along the proximal-distal axis were classified as “Rotate” cells. The solid lines show the model’s best fits, and the shaded regions indicate the 95% confidence intervals. Complementary to the proportion data in (B), in Y and AU rats, the probability of being a “Rotate” cell increases from proximal to distal CA3. In contrast, in AI rats, the probability of being a “Rotate” cell decreases from proximal to distal CA3.

There were notable group differences across the CA3 subregions in the proportion of the response types (Fig. 2B, Supp. Table 1). In Y rats, there was a decreasing proportion of Remap cells, with a correspondingly increasing proportion of Rotate cells, from proximal to distal CA3. This trend is consistent with previous findings in Y rats reported by Lee et al. (2015). AU rats showed a similar trend as Y rats, although the proportions in each region were different from Y rats. However, the trend was opposite in AI rats, with a decreasing proportion of Rotate cells and an increasing proportion of Remap cells from proximal to distal CA3. Specifically, in the Y rats, a high proportion (76%) of cells in proximal CA3 remapped (n = 99/130), compared to 65% of cells in intermediate CA3 (n = 75/115) and 55% of cells in distal CA3 (n = 31/56). In AU rats, 57% of cells in proximal CA3 remapped (n = 81/142), compared to 48% in intermediate CA3 (n = 52/109), and 39% in distal CA3 (n = 18/46). In contrast, in AI rats, only 28% of cells remapped in proximal CA3 (n = 24/85), 35% in intermediate CA3 (n = 58/165), and 43% in distal CA3 (n = 20/47). There was a significant association between the proportion of response types and groups in the proximal (*X*^2^_(2,357)_ = 48.19, p < 0.0001) and intermediate (*X*^2^_(2,389)_ = 24.56, p < 0.0001) regions but not in the distal region (*X*^2^_(2,149)_ = 3.06, p = 0.22), suggesting response types differ among the groups in proximal and intermediate CA3 but not in distal CA3.

Even though the CA3 transverse axis was historically divided into 3 discrete regions (CA3a, b, and c) (Lorente de No, 1934), there are no defining criteria for discrete borders, and anatomical (Ishizuka et al., 1990; Li et al., 1994; Witter, 2007) and genetic (Lein et al., 2007; Thompson et al., 2008) studies suggest that the axis is more of a continuum displaying a functional gradient (Hunsaker et al., 2008; Lee et al., 2015; Lu et al., 2015; Marrone et al., 2014; Nakamura et al., 2013). Thus, we next performed a logistic regression analysis to model the probability of a cell being a “Rotate” cell along the transverse axis, measured as a continuous variable (Fig. 2C). In AI rats, the probability of being a “Rotate” cell was high towards the proximal region but decreased towards the distal region. In contrast, in Y and AU rats, the probability of being a “Rotate” cell was low towards the proximal region but increased towards the distal region. Statistical analyses on the interaction between groups and distance along the transverse axis showed that the average curves across the groups differed significantly between Y and AI (p = 0.0002) and between AU and AI (p = 0.02) but the difference between Y and AU rats showed only a trend (p = 0.07). Thus, consistent with the within-region comparisons shown in Fig. 2B, Y and AU rats have an opposite trend compared to AI rats in the proportion of cell types along the transverse axis, with AU rats showing results that appear intermediate between the Y and AI groups.

### Population responses in the cue-conflict manipulations along the CA3 transverse axis

The previous analysis is useful to show, at the level of single place fields, how CA3 responds differentially across the groups and locations along the transverse axis. However, that analysis did not allow for investigation of ensemble coherence and the degree of potential rate remapping that might occur. To address these issues, spatial correlation matrices from the population firing rate vectors at each location on the circular track were analyzed (Lee et al., 2015, 2004; Neunuebel and Knierim, 2014) (Fig. 3A). To create these matrices, the firing rate of each cell was calculated for each 1° bin on the track and normalized to its peak rate. The firing rate maps of all *n* cells in the sample were stacked to create a 360 x *n* matrix, in which each column of the matrix represents the population firing rate vector for each angle of the track (Supp. Fig. S2). The firing rate vectors at each angle of a STD session (STD1) were correlated with the firing rate vectors at each angle of the next STD session (STD2), to create STD1 x STD2 correlation matrices, or with the next MIS session, to create STD1 x MIS correlation matrices. To make statistical comparisons across the groups and CA3 subregions tractable, we collapsed the data from all 3 rotation mismatch amounts. Because prior studies of CA3 with this same cue-mismatch manipulation have shown that place fields tended to be controlled by the local cues (i.e., the place fields rotated along the track by increasing amounts when the local cues were rotated by increasing amounts; Supp Fig. S3) (Lee et al., 2015, 2004; Neunuebel and Knierim, 2014), we first adjusted the firing rate maps to compensate for the different degrees of cue rotation. To do so, the mean rotation angle of place fields for all recorded cells of a MIS amount (90°, 135°, or 180°) was calculated (Supp. Fig. S2). Each cell’s firing rate map in each MIS session was then rotated by this mean rotation amount. Thus, if the place fields rotated coherently in the MIS sessions, controlled by one set of cues, rotating the rate maps in the MIS session by the mean rotation amount would align the location of the place fields in MIS sessions with the location of the place fields in the preceding STD1 session. This alignment would create a strong correlation band along the main diagonal in the spatial correlation matrices between the STD and MIS sessions. However, if the cells did not rotate coherently in the MIS sessions or if the cells remapped, rotating the place fields in the MIS session by the mean rotation amount would misalign the locations of the place fields in MIS and STD sessions, creating a dispersed and weak correlation band along the main diagonal in the spatial correlation matrices between the STD and MIS sessions.

**Figure 3.**
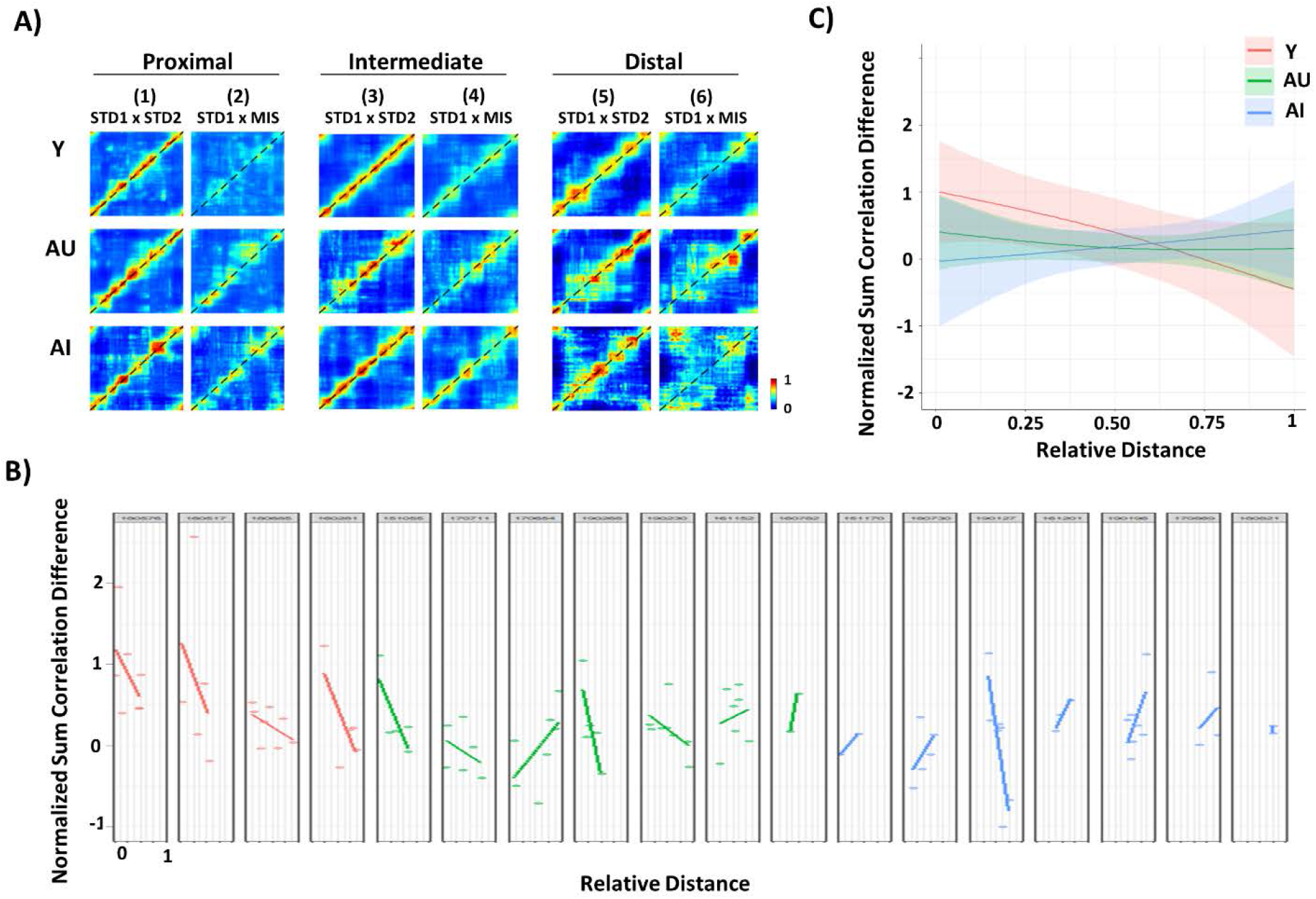
Correlated representations between the STD and MIS sessions differ in age groups along the transverse axis. **(A)** STD1 x STD2 matrices (columns 1, 3, 5) and STD1 x MIS matrices (columns 2, 4, 6) in each CA3 subregion for all groups. **(B)** The correlation differences along the transverse axis organized according to each animal. Each dot represents a tetrode recorded in that region along the axis. **(C)** A linear mixed effects model was used to compare the correlation differences along the transverse axis as a continuous variable. The solid lines show the model’s best cubic-splines fits, and the shaded regions indicate the 95% confidence intervals. There were differences in the shapes of the fitted curves across the animal groups. There was a significant difference in correlation difference between Y and AI groups but not between Y and AU or AU and AI groups.

Figure 3A demonstrates the population coherence between repeated sessions in the STD sessions and between the STD and MIS sessions. Columns 1, 3, and 5 show correlation matrices for STD1 vs. STD2 sessions for all groups and CA3 regions. In each case, there is a band of high correlation along the main diagonal of the matrix (dashed line), indicating that the population representation of spatial location on the track was similar (i.e., coherent) between the two STD sessions (see also Fig. S3). Columns 2, 4, and 6 show correlation matrices between the STD and MIS sessions. For Y rats, the correlation band along the main diagonal of proximal CA3 was very weak, indicating that there was a large amount of spatial remapping between the two sessions. For intermediate and distal CA3, however, the correlation band was stronger, replicating the result of Lee et al. (2015) of a gradient from pattern separation to pattern completion along the transverse axis of CA3. Both AU and AI rats showed stronger proximal CA3 correlation bands than Y rats, and less of a difference in the strength of the bands from proximal to distal CA3.

The total correlation along the principal diagonal of the matrix measures the degree of coherence in the firing of cells across the two experimental sessions. Our hypotheses about the effects of region and group on the degree of coherence can therefore be tested by comparing the sum of the diagonal correlations among different regions and groups of animals. Although the firing locations of all place cells between the STD1 and STD2 sessions were stable across all groups (Supp. Fig. S4), we note that the different groups and CA3 subregions may have quantitatively different degrees of coherence or correlation in their spatial maps across sessions due to differences in place field size, firing rates, and other factors that might affect the values in the correlation matrices (compare STD1 vs. STD2 matrices in Fig. 3A). To control for such inherent variability, we calculated the total correlation *differences* between the main diagonals of the STD matrix (STD1 x STD2) and the MIS matrix (STD1 x MIS matrix) (Supp. Fig. S2) to isolate the correlation differences that arise from the mismatch manipulation rather than the inherent place field differences. Thus, if the correlation strength in the STD matrix is stronger than the correlation strength in the MIS matrix, the correlation difference will be more positive. If the correlation strengths between the two matrices are similar, the correlation difference will be nearer 0. To examine effects of age and cognitive status on a continuum across the CA3 transverse axis, population correlation matrices for STD1 vs. STD2 and STD1 vs. MIS sessions were calculated for each tetrode, similar to that of Fig. 3A but restricted to the population of cells recorded on that tetrode. The total correlation differences between the STD1 vs. STD2 matrix and the STD1 vs. MIS matrix for each tetrode along the transverse axis for each animal are shown in Fig. 3B. Notably, all (4/4) Y rats showed a decreasing trend along the transverse axis, while 4/7 AU rats and only 1/6 AI rats showed a decreasing trend (one AI rat did not have a spread of tetrode locations along the transverse axis and thus no slope could be assessed for that rat). There are not enough data points to run a valid chi-squared test on these numbers, but the trends appear to indicate, on a per-animal basis, a shift from decreasing pattern separation along the transverse axis (from proximal to distal) in Y rats to increasing pattern separation along the axis in AI rats. AU rats are intermediate between the Y and AI rats.

To statistically analyze the trends across animals, we employed linear mixed effects models (Laird and Ware, 1982), which take into account inhomogeneities across animals in sampling along the transverse axis as well as correlations among cells within an animal. Because the aged rats tend to run at a slower speed than Y rats and speed can be a confounding variable that may contribute to place cell firing rates (Lee et al., 2021), momentary speed was incorporated as a fixed effect. There was a statistically significant difference in the shape of the fitted curves between Y and AI rats (p = 0.04), but not between Y and AU rats (p = 0.22) or between AU and AI groups (p = 0.57) (Fig 3C). From these tests (Figs. 2-3), we can conclude that the normal gradient from less correlation (i.e., pattern separation) to high correlation (i.e., pattern completion) observed along the transverse axis in Y rats is abnormal in AI rats, with AU rats showing an intermediate pattern between the Y and AI rats.

### Global-cue rotation only in CA3

Since the MIS sessions require recognition of a mismatch between the global and the local cues, a control experiment was performed to verify that aged rats were able to see the global cues. The textured track was replaced with a solid wooden track to remove the salient, local texture cues on the track. All rats were then given STD1-MIS-STD2 sessions where only the global cues were rotated by 45° or 90° in the clockwise direction in the MIS session. Population coherence demonstrated global cue control when local cues were removed (Supp. Fig. S5), suggesting that the aged rats could perceive the global cues and that the place cells can be controlled by the global cues in the absence of conflicting local-cue information (Knierim and Hamilton, 2011).

### Two-shape experiment

To test whether the interaction between the groups and CA3 subregions seen in the cue-mismatch experiment generalized to other conditions of remapping, a subset of rats from the cue-mismatch experiment were recorded during open-field foraging in two different shapes, a square (S) and a circle (C) enclosure, placed at a constant location in room A. Three Y, 6 AU, and 6 AI rats were recorded for two days with the session sequence as either S-C-C-S or C-S-S-C. Example cells from Y rats showed remapping between the two shapes (i.e., fired only in one shape) in proximal and intermediate CA3 but fired in both shapes in distal CA3 (Fig. 4). Example cells from AU rats showed remapping in proximal CA3 but fired in both shapes in intermediate and distal CA3. In contrast, example cells from AI rats fired in both shapes in all CA3 subregions.

**Figure 4.**
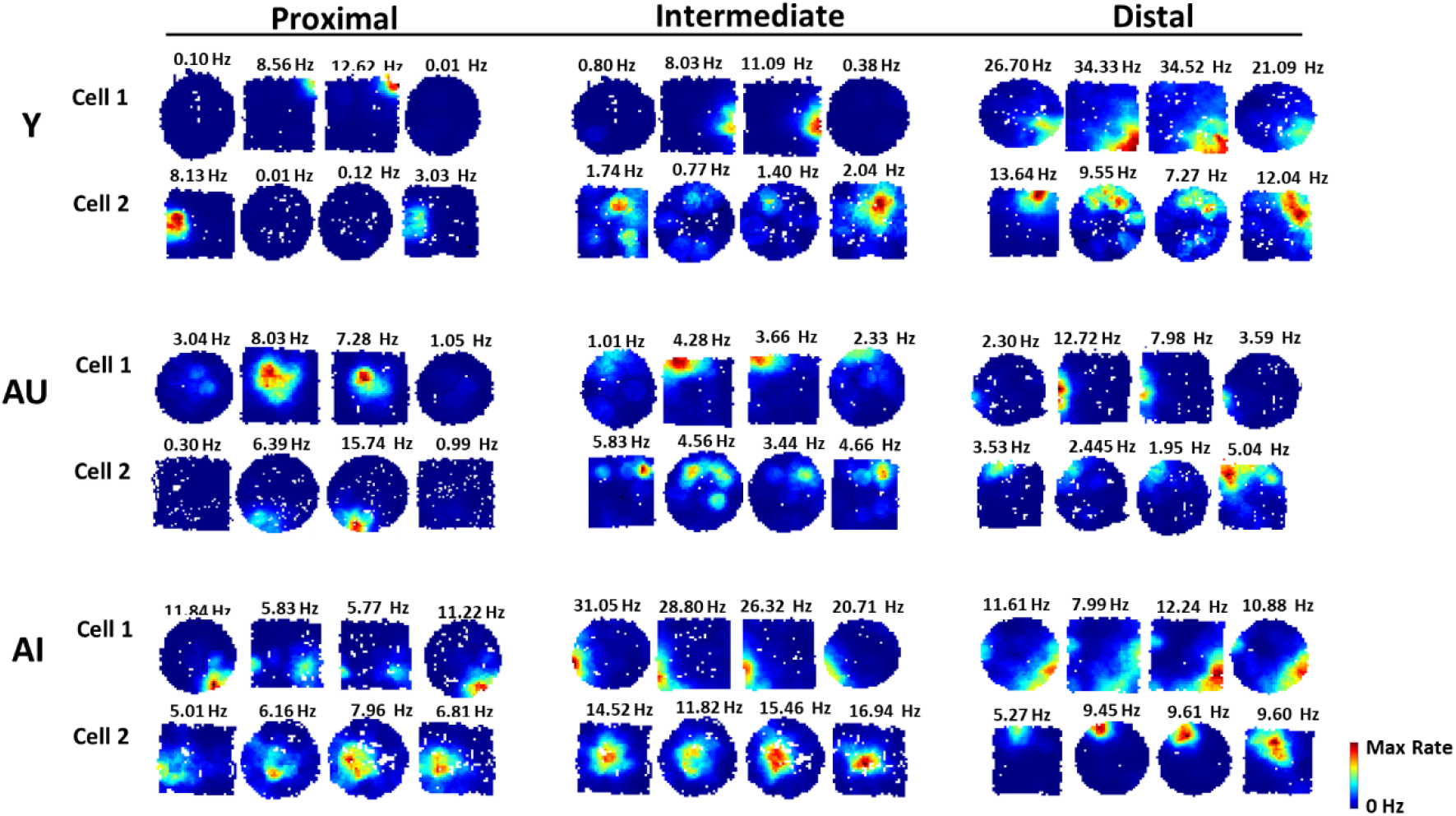
Representative rate maps for Y, AU and AI rats showing different degrees of remapping in the two-shape experiment. Rate maps of two example cells from each age group in each CA3 subregion. Example cells in Y rats showed remapping between the two shapes in proximal and intermediate CA3 but fired in both shapes, with similar place fields, in distal CA3. Example cells in AU rats showed remapping in proximal CA3, but fired in both shapes, with similar place fields, in intermediate and distal CA3. Example cells in AI rats fired in both shapes, with similar place fields, in all CA3 subregions. These representative cells have spatial correlation values between the different shapes that are within the second quartile or the third quartile from the population distribution.

To quantitatively assess the degree of remapping between the two shapes, we calculated population vector (PV) correlations (Fig. 5A), which are sensitive to the distribution of changes in the cells’ firing locations and firing rates (i.e., global and rate remapping) (Leutgeb et al., 2005). Of the 3 Y rats recorded in the two-shape experiment, 1 anomalous Y rat showed remapping between the same shapes (i.e., the place field locations changed between the visits to the same shapes). The repeated visits to the same environment are meant to serve as a control for the magnitude of correlation between the repeated visits to the same environment, against which we can compare the amount of correlation (remapping) between visits to the different environments. This is done by subtracting the correlation in the different shape from the correlation in the same shape. It is highly unusual for a Y rat to show remapping between visits to the same environment under normal circumstances. Since the anomalous rat showed remapping in the same shape, the same shape can no longer be considered a control that serves as a baseline measure of correlation without remapping (Supp.Fig. S6). Thus, this anomalous rat was excluded from the analysis (Supp. Table 2). The data in the two-shape condition was from the remaining 2 Y rats, but the results observed in Y rats are consistent with prior published studies showing stable firing of CA3 cells in the same shape (Fyhn et al., 2007; Leutgeb et al., 2007, 2005), thus lending credence to the results seen in the Y rats.

**Figure 5.**
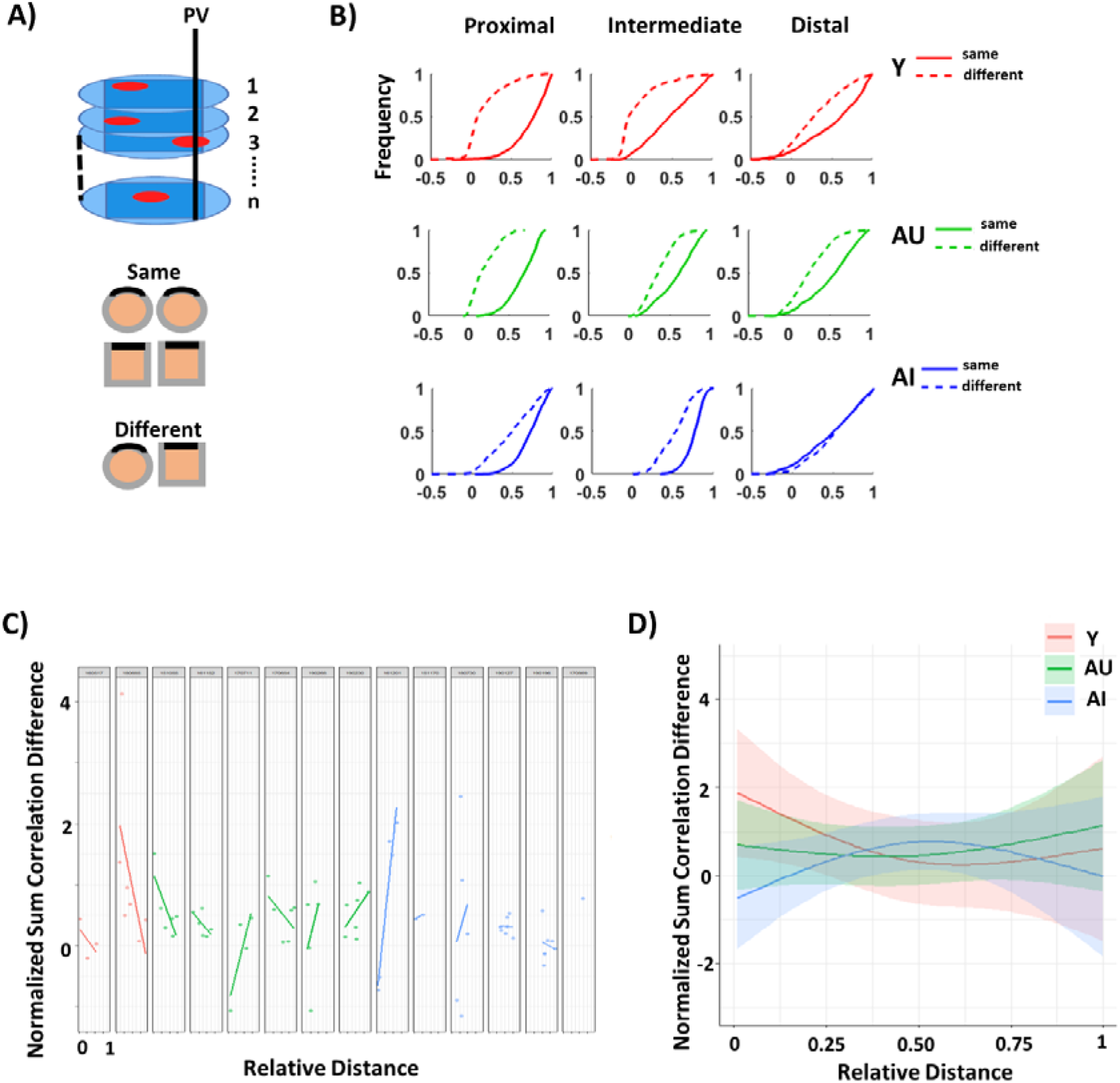
Representations for the different shape environments are less distinguishable in AI rats compared to Y and AU rats. **(A)** Illustration showing the population vector correlation analysis (top). The rate maps of all CA3 cells were stacked, and the firing rates along the z axis represent the population vector (PV) for each x-y bin that was shared between the circle and the square. (Unshared bins between the two shapes were excluded). Correlations between the population vectors were computed between the same shapes (C-C or S-S) and the different shapes (C-S). **(B)** Cumulative distribution plots for PV correlations between the same shape (solid lines) and the different shape (dashed lines) in Y (top row), AU (middle row), and AI rats (bottom row). PV correlation of 0 indicates strong remapping and PV correlation of 1 indicates completely correlated firing patterns between the two environments. **(C)** Observed PV correlation differences along the transverse axis organized according to each animal. Each dot represents a tetrode recorded at that location along the axis. **(D)** The PV correlation difference for each tetrode location across all animals. The solid lines show the model’s best cubic-spline fits, and the shaded regions indicate the 95% confidence intervals. There were differences in the shape of the fitted curves between Y and AI groups but not between Y and AU or between AU and AI groups.

In all groups, population vectors in the same shape configurations (Fig. 5B, solid lines) were more strongly correlated than the population vectors in the different shape configurations (dashed lines). However, there were noticeable differences in the magnitude of PV correlation strengths between the shape configurations. As we did in Figure 2B and Fig. 3A, we first analyzed data according to the classic definitions of proximal, intermediate, and distal CA3 subregions, for visual illustration. In Y rats, the change in PV correlations in the same shape versus the different shape was greater in proximal and intermediate CA3 than in distal CA3. In AU rats, the change in PV correlations was greater in proximal CA3 than in intermediate and distal CA3. In AI rats, the change in PV correlations was small in all CA3 subregions.

To measure the change in PV correlation strengths between the same shapes and the different shapes on a continuum across the CA3 transverse axis, the PV correlation change was calculated for the population of cells recorded on each tetrode. First, we summed the PV correlations for each of the bins (i.e., each of the values in the cumulative distribution plots as in Fig. 5B). We then subtracted the summed PV correlation for the different shapes from the summed PV correlation for the same shapes, and divided this difference by the sum of the two quantities. The PV correlation difference along the transverse axis organized according to each animal showed heterogeneous relationships across the groups (Fig. 5C).

A linear mixed effects model that incorporates momentary speed as a fixed effect was used to compare the PV correlation strengths between the groups and the location along the transverse axis as a continuous variable. As in Figures 2C and 3C, the model took into account heterogeneities in sampling across individuals and groups, as well as correlated firing within animals. There were differences in the shape of the fitted curves across the groups (Fig. 5D). There was a significant difference between Y and AI groups (p = 0.03), but not between Y and AU groups (p = 0.36) or between AU and AI groups (p = 0.14). Similar to the results observed in Fig. 3C, the gradient from less correlation (i.e., pattern separation) to high correlation (i.e., pattern completion) observed along the transverse axis in Y rats was not observed in AI rats.

### Responses in the two-room experiment

For the two-room experiment, after session 4 in room A, the same rats from the two-shape experiment were taken to room B and were given additional sessions (sessions 5 and 6) in the square and the circle. All comparisons were made between session 4 in room A (last session in room A) and session 5 or 6 in room B (whichever session matched the same shape as session 4 in room A). Example rate maps showed that place fields changed their firing locations between the two rooms, or were active in only one room, in all CA3 subregions in Y, AU, and AI rats (Fig. 6A). Population vector correlations were very low (Fig. 6B) in all CA3 subregions for all groups. The sum of the PV correlations for the population of cells recorded on each tetrode along the transverse axis was calculated and compared across the groups using a linear mixed effects model that incorporates momentary speed as a fixed effect. There was no difference in the shape of the fitted curves across the groups [between Y and AU (p = 0.90), between Y and AI (p = 0.85), and between AU and AI (p = 0.86)] (Fig. 6C). These data suggest that when spatial location changes between two environments with distinctive features, all groups, irrespective of age and cognitive status, are able to dissociate the different rooms by remapping the place fields in all CA3 subregions.

**Figure 6.**
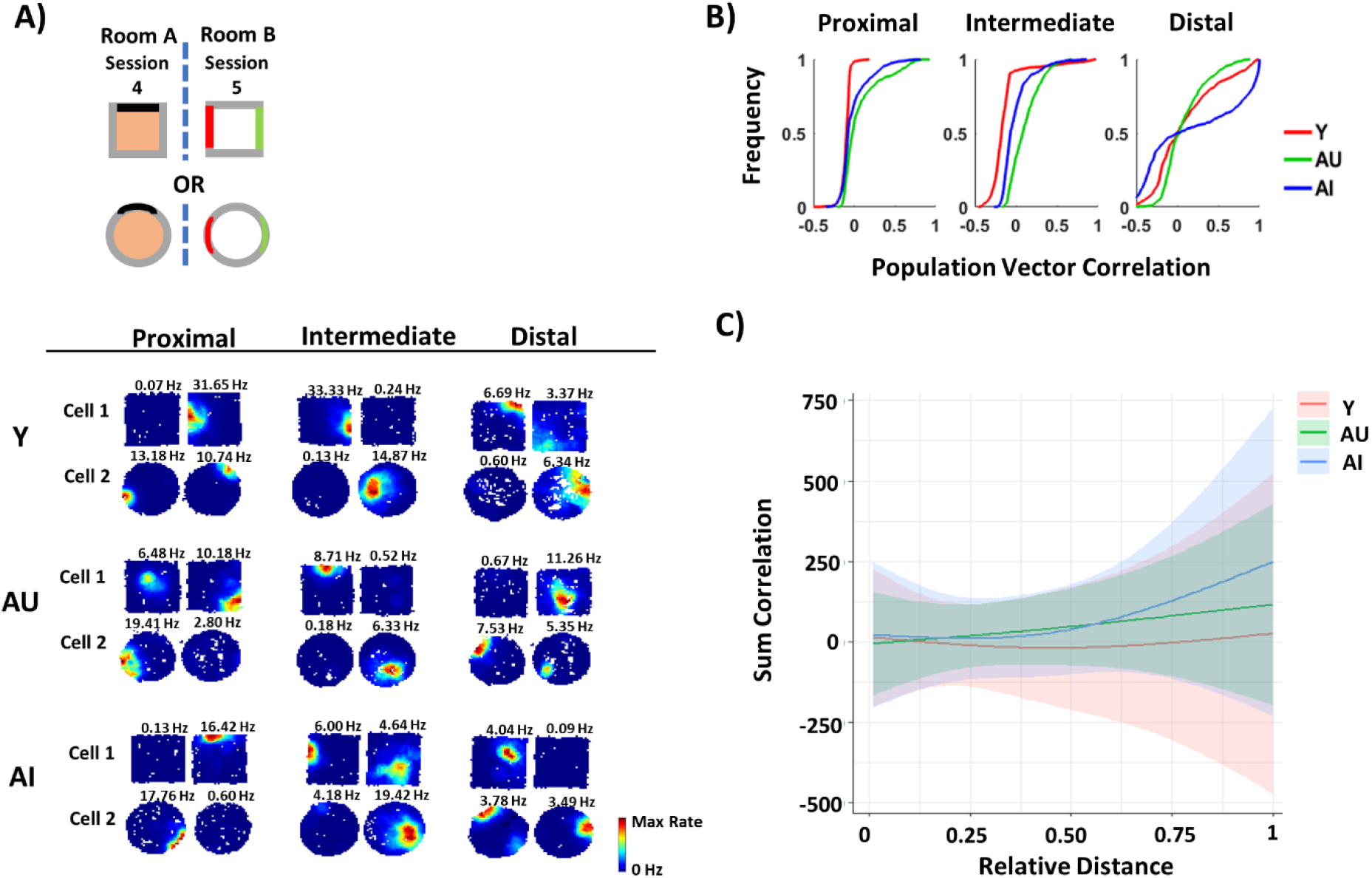
CA3 place cells in all age groups remap between the two rooms. **(A)** Schematics showing the two-room task (left). Rate maps of two example cells from each animal group in each CA3 subregion. Cells show remapping between the two rooms in all CA3 subregions in all age groups. **(B)** Cumulative distribution plots for PV correlation between Room A and Room B in each CA3 subregion. **(C)** The PV correlation difference for each tetrode location across all animals. The solid lines show the model’s best cubic-spline fits, and the shaded regions indicate the 95% confidence intervals. There were no differences in the shape of the fitted curves among the age groups.

## Discussion

The current study showed a shift from pattern separation to pattern completion along the CA3 transverse axis in Y rats that was not observed in AI rats. In both the cue-mismatch and the two-shape experiments, Y rats showed a decrease in remapping from proximal to distal CA3, while AI rats showed strong correlation in all CA3 subregions. The trends in AU rats were in between Y and AI rats, with no significant difference compared to either Y or AI rats. Results from these two experiments suggest that in Y rats, there is a functional transition along the CA3 transverse axis, with a bias toward pattern separation in proximal CA3 (i.e., more remapping) and a bias toward pattern completion in distal CA3 (i.e., less remapping). This functional shift was not evident in AI rats, as the results suggest pattern completion in all CA3 subregions in AI rats. In contrast, in the two-room experiment, there was remapping in all CA3 subregions in all groups, suggesting that when the two environments do not share overlapping features and are in distinctly different locations (i.e., two different rooms), CA3 cells in all subregions discriminate between the two environments regardless of age and cognitive status. In agreement with prior studies on age-related hyperactivity of proximal CA3 cells (Lee et al., 2021), these data suggest that the functional impairment along the CA3 transverse axis is observed towards the proximal region rather than the distal region.

### Pattern separation and pattern completion along the CA3 transverse axis in aging

Prior studies of neurocognitive aging in the model used here suggested that enhanced pattern completion might be favored in aged rats as the result of the hyperactivity in the pattern completion network of CA3 (Wilson et al., 2005, 2006). Based on the functional dissociation along the CA3 transverse axis, the simple logic is that hyperactivity in aged rats should be observed in distal CA3, which supports pattern completion. However, distal CA3 does not show hyperactivity in aged rats, and may even show hypoactivity; instead, proximal CA3 shows hyperactivity (Lee et al., 2021). Since proximal CA3 supports pattern separation (Lee et al., 2015; Lu et al., 2015), how does hyperactive proximal CA3 favor pattern completion instead of pattern separation in aging? To resolve this apparent paradox, we previously proposed a model (Fig. 7A) of how age-related changes in CA3 firing rates along the transverse axis affect putative attractor dynamics of CA3 (Lee et al., 2021). We proposed that because proximal CA3 in aged rats presumably receives disrupted pattern-separated outputs from DG, the recurrent collaterals in proximal CA3, facilitated by the hyperactivity, may override the DG inputs to produce a single attractor basin that results in pattern completion. The possible hypoactivity in distal CA3 may result in weaker attractor basins in aged rats to result in bistable representations of the same environment, such as have been observed under certain conditions in aged rats (Barnes et al., 1997; Rapp, 1998; Redish et al., 1998; Tanila, 1998). The current study is consistent with the model by providing direct physiological evidence that representations in AI rats show abnormally high pattern completion towards the proximal end of CA3. The difference between Y and AI rats may be a prime example of the more general phenomenon of age-related dedifferentiation, which refers to a reduction in functional selectivity of brain regions with increasing age (Koen and Rugg, 2019; Koen et al., 2020). The present results suggest that there is such a dedifferentiation of functional specificity within a brain area, as the normal differentiation of function along the CA3 axis of Y rats transition to a more homogeneous functionality along this axis in AI rats.

**Figure 7.**
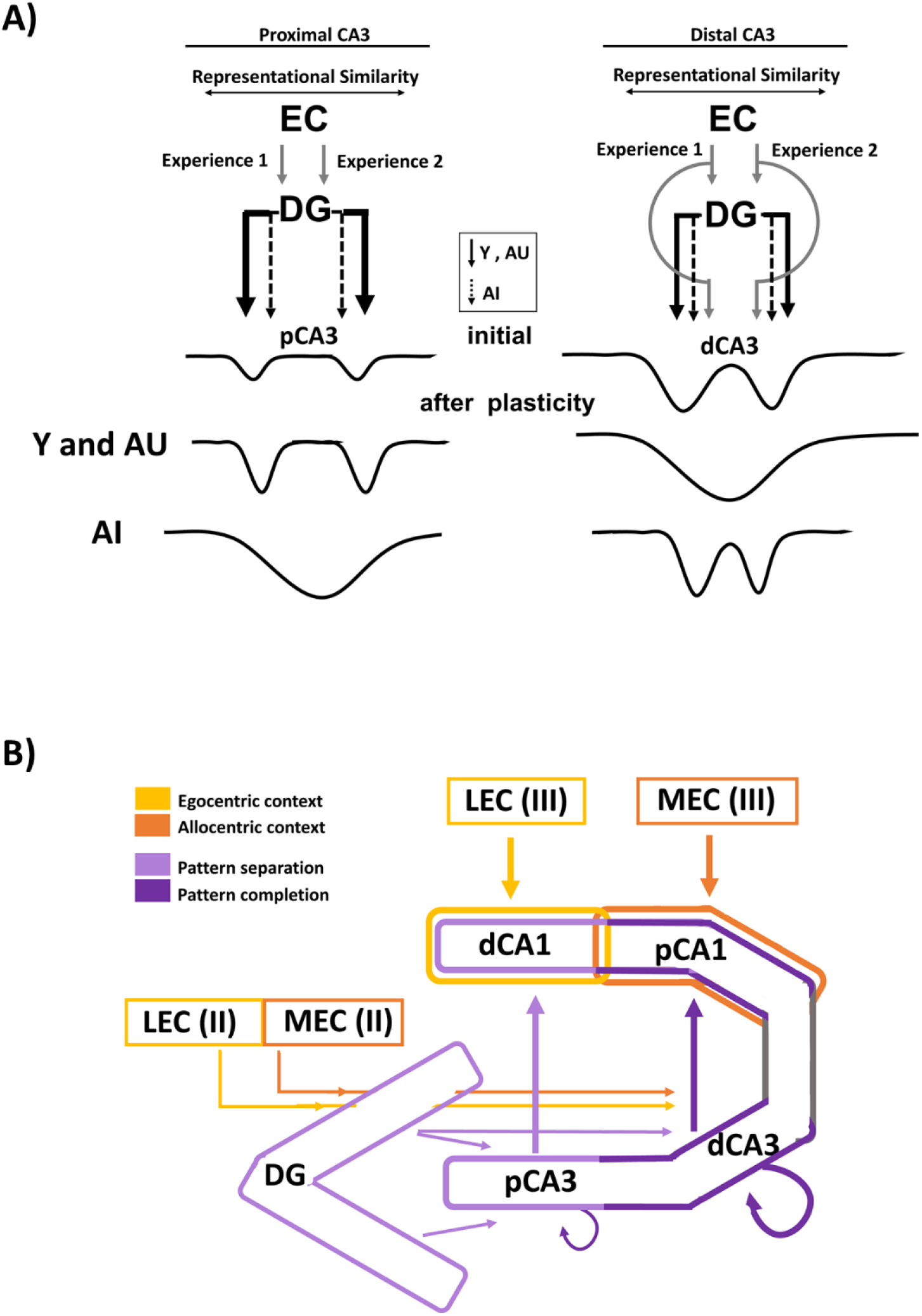
Hypothesized model of how the attractor dynamics of CA3 is affected by aging. **(A)** In proximal CA3 (left), the EC neural representations of similar experiences (shown as distance between two input arrows along a “representational similarity” axis) will be pattern separated in the DG (indicated by the larger distance between the DG output to CA3 than the distance between the two EC inputs shown in thick, black solid lines). Proximal CA3, which receives stronger inputs from the DG (both upper and lower blades) coupled with minimal EC inputs, imposes the separated patterns on the CA3 attractor network. Through learning, the recurrent collaterals of pCA3 increase the strength of the attractor basins, resulting in pattern separation in pCA3 for Y and AU rats. In contrast, for AI rats, dysfunction of the EC inputs and the local inhibitory circuitry of the DG hilus are hypothesized to reduce the ability of the DG to separate the similar EC input patterns (indicated by the close distance between the DG outputs to CA3 shown in black, broken lines). The hyperactivity of pCA3 cells in AI rats may further increase the relative strength of the pCA3 to override the DG inputs and cause a single basin of attraction, resulting in pattern completion in pCA3. In distal CA3 (right), there are weaker DG inputs (black solid lines) and stronger EC inputs (gray lines) compared to pCA3. The combined DG and EC inputs may impose initial patterns on dCA3 that are somewhat closer together and overlapping than those imposed in pCA3, and the stronger recurrent collateral system of dCA3 may merge the two attractor basins for a broader and deeper basin, resulting in pattern completion for Y and AU rats. In AI rats, weaker DG inputs (black, broken lines) and the possible hypoactivity of dCA3 cells may result in weaker attractor basins in dCA3 and prevent the merging of the attractor basins, resulting in two attractor states with relatively low energy barriers between them. (modified from Lee et al., 2021) **(B)** Parallel processing streams within the hippocampus. Allocentric contextual information (dark orange) from MEC projects to distal CA1 (dCA1) and egocentric content information (yellow) from LEC projects to proximal CA1 (pCA1). Both MEC and LEC directly project to the DG as well as to CA3, with stronger projection to distal CA3 (dCA3). The DG projects to CA3, with stronger projection to proximal CA3 (pCA3). The DG and pCA3, involved in pattern separation, are marked in light purple, and dCA3, with its strong recurrent circuitry, involved in pattern completion is marked in dark purple. Pattern-separated outputs from pCA3 preferentially project to dCA1, while pattern-completed outputs from dCA3 preferentially project to pCA1.

In addition to pattern separation vs. pattern completion, several mnemonic functions have been proposed for the DG-CA3 network, including contextual discrimination during memory encoding and recall (Hainmueller and Bartos, 2020), the binding of objects and events to spatial contexts (Lee and Jung, 2017) and sequence encoding and recall (Lisman, 1999; Lisman et al., 2005). While it is important to determine the exact function of each hippocampal subregion in mnemonic processing, these theories may not be mutually exclusive, as they all attempt to capture how the different parts of the hippocampus and related circuitry contribute to encoding, storage, and recall of memories. Further experiments designed to test predictions from these models, and how these predictions may vary depending on location along the CA3 transverse axis and on cognitive status in aging, will illuminate the precise functions of the complex DG-CA3 circuit.

### Implications for aging

What are the implications of age-related deficits in proximal CA3? Previous studies have argued that proximal CA3 should be considered more as a functional component of the DG pattern-separation circuit than as a functional component of the distal CA3 pattern completion circuit (Hunsaker et al., 2008; Lee et al., 2015, 2020, 2021). The outputs from proximal and distal CA3 are topographically organized along the CA1 transverse axis, forming parallel processing streams along the hippocampal transverse axis (Fig. 7B). Pattern-separated output from proximal CA3 and egocentric sensory information from lateral entorhinal cortex (LEC) target distal CA1 while pattern-completed output from distal CA3 and allocentric spatial information from medial entorhinal cortex (MEC) target proximal CA1 (Amaral and Witter, 1989; Ishizuka et al., 1995; Lee et al., 2020). In aging, there is compelling evidence that the LEC is selectively vulnerable, including evidence of neurofibrillary tangles in the LEC earlier than the MEC in human aging (Braak and Braak, 1995); hypometabolism in the LEC, but not in the MEC, in patients with mild cognitive impairment (Khan et al., 2014); and significantly reduced expression of reelin, a protein involved in cell migration and synaptic plasticity, in layer II of LEC, but not in MEC, of AI rats compared to Y and AU rats (Stranahan et al., 2011a, 2011b). Profound impairment in the pattern separating function of the DG/proximal CA3 and the functions of the LEC suggest that the pCA3-dCA1-LEC pathway is more vulnerable in aging than the dCA3-pCA1-MEC pathway. MEC appears to be more associated with path integration and the creation of allocentric maps (the spatial *context* of experience), whereas LEC appears to be more associated with egocentric representations of the specific *content* (and perhaps temporal context; (Tsao et al., 2018)) of an experience (Knierim, 2015). Distal CA1, with its specific, reciprocal connectivity with LEC, appears to require input from proximal CA3 that is biased toward pattern separation to perform its function, whereas proximal CA1, with its specific, reciprocal connectivity with MEC, appears to require input from distal CA3 that is biased toward pattern completion. It is not known why these two processing streams require differential balances between pattern completion and pattern separation from CA3, but it is possible that the MEC-related pathway is biased toward pattern completing small changes to an environment, in order to retrieve the appropriate spatial context, and the LEC-related pathway is biased toward pattern separating representations of the specific items that were altered, in order to unambiguously encode similar events that occur within the same context. In this scenario, the mnemonic deficits associated with aging arise more from dysfunction in the pattern separation processing of the individual items of experience, rather than the ability to correct for small errors or degraded inputs that arise from small changes to the representations of the allocentric, spatial contexts within which experiences occur.

Consistent with studies of aged animals, fMRI studies in aged individuals with deficits in lure discrimination tasks show heightened activity in DG/CA3 (Bakker et al., 2012, 2015; Reagh et al., 2018; Yassa et al., 2011). While it is difficult in human fMRI studies to distinguish DG BOLD signals from CA3 signals, it is possible that the DG/CA3 heightened activity in BOLD signals may largely reflect the heightened activity in proximal CA3, as proximal CA3 appears to be disproportionately large in humans, with ∼75% of CA3 apparently homologous to rodent proximal CA3 (Lim et al., 1997). Accordingly, the proportional increase in the size of proximal CA3 from rodents to primates suggests that adaptive pressures to support human episodic memory increased the relative role of the pCA3-dCA1-LEC pathway compared to the dCA3-pCA1-MEC pathway, and disruption to the former pathway results in the age-related deficits in episodic memory.

### Cognitive resilience in AU rats

While memory function is vulnerable to aging, a subpopulation of aged animals (including aged humans) is able to retain normal memory function. We previously showed that hyperactivity localized towards proximal CA3 region was present in both AU and AI rats (Lee et al., 2021). While AU rats did not show statistical difference from AI rats, they also did not show statistical difference from Y rats, suggesting that AU rats are not as impaired as AI rats. Numerous examples in the neurological literature demonstrate gradual decreases in neural properties with little behavioral deficit until a threshold is crossed, after which impaired behavior/cognition becomes apparent (e.g., Parkinson’s Disease and Alzheimer’s Disease (Engelender and Isacson, 2017; Palasí et al., 2015)). Although aged rats were grouped into impaired and unimpaired categories for analysis purposes, in reality the performance of these rats in the water maze lies on a continuum, ranging from Learning Index (LI) scores as impressive as the best Y rats, through LI scores just outside the range of normal scores of Y rats, to scores that are many standard deviations worse than Y rats (Gallagher et al., 1993). There is a relatively continuous gradient in the physiological results of our study as well, as AU rats tend to show results intermediate between Y and AI (Fig. 2, 3, 5). It appears that AU rats may be physiologically on their way to behavioral impairment, but have not crossed a threshold yet. In addition, compensatory adaptive mechanisms in aging may allow them to perform well even as alterations in their brains are becoming more dysfunctional compared to Y. The level of their physiological impairment may not be great enough to show a statistically significant difference from Y rats or from AI rats, but the trends in the data are clear and consistent.

There are distinctive changes in hippocampal circuitry that may account for compensatory mechanisms in AU rats. Postsynaptic inhibition is enhanced in granule cells in AU rats compared to AI rats (Tran et al., 2018, 2019), and somatostatin-positive HIPP interneurons are intact in AU rats but reduced in AI rats (Spiegel et al., 2013). Enhanced inhibition in granule cells in AU rats may support sparse coding in the DG, which is essential for pattern separation (Koh et al., 2020). With respect to the evidence that aging has a greater effect on the role of the pCA3-dCA1-LEC pathway compared to the dCA3-pCA1-MEC pathway, the loss of reelin expression in LEC neurons occurs in the condition of memory impairment in both AI rats and rhesus monkeys, with relative sparing of that condition in AU animals in both species (Long et al., 2020; Stranahan et al., 2011a, 2011b). There is also a redistribution of synaptic weights between AI and AU age groups in rodents, where AMPAR expression in stratum lucidum (the layer where mossy fibers make synaptic contacts with CA3) is higher in AU rats but AMPAR expression is significantly higher in stratum radiatum (the layer where CA3 makes autoassociational connections) in AI rats (Buss et al., 2021). Enhanced AMPAR expression in stratum lucidum may support increased feedforward activation of pattern separated outputs to CA3. However, exactly how these changes are controlled and provide compensatory mechanisms in AU rats still remains largely unknown and needs further study.

### Summary

The present study demonstrates that age-related impairments in pattern separation is localized preferentially towards the proximal CA3 in AI rats. When changes are made to the environment, neural representations in Y rats are more orthogonal in proximal CA3 than in distal CA3, suggesting a functional gradient (pattern separation to pattern completion) along the CA3 transverse axis. In contrast, AI rats show less orthogonal representations in proximal CA3 compared to Y rats but more generalized representations in distal CA3 that are similar to Y rats. Thus, enhanced pattern completion in AI rats may result from proximal CA3 that is more biased towards pattern completion rather than pattern separation. Furthermore, AU rats show trends that are intermediate to Y and AI rats, suggesting that AU rats are not equivalent to young adults.

## Methods

### Subjects and Surgery

Male Long-Evans rats (retired breeders at 9 months) were obtained from Charles River Laboratories (Raleigh, NC) and housed in a vivarium at Johns Hopkins University until age 22-26 months. Young rats (3-6 months) were also obtained from Charles River Laboratories. Prior to the onset of water maze behavioral testing, rats were individually housed with ad libitum access to food and water during a 12-hour light/dark cycle. After the water maze testing, rats underwent hyperdrive implant surgery and were housed during a reversed 12-hour light/dark cycle. All recordings were done in the dark cycle. All animal care, housing, and surgical procedures conformed to the National Institute of Health standards using protocols approved by the Institutional Animal Care and Use Committee at Johns Hopkins University.

All rats were prescreened for spatial learning ability in the Morris water maze as previously described (Lee et al., 2021) prior to the implantation of recording electrodes. In total, 14 aged rats were tested at 22-24 months and 4 young rats were tested at 3-4 months in the water maze. These rats were trained for 8 days (3 trials per day) to locate a submerged escape platform that remained at the same location in the water maze tank, with every sixth trial as a probe trial (no escape platform for the first 30 seconds). The learning index (LI), an average of weighted search proximity scores obtained during probe trials (Gallagher et al., 1993), was used to classify animals as aged-impaired (AI; LI > 240) or aged-unimpaired (AU; LI < 240), using an a priori LI criterion from previous studies. A LI of 240 represents 2 S.D. above the mean of a large population of Y animals (Gallagher et al., 1993). On day 9, rats were given six trials to locate a visible platform above the water level to screen for nonspecific task impairment such as swimming behavior and escape motivation. Following water maze testing, rats were placed on restricted food access to reduce their body weight to 85% while they were given foraging sessions (20 mins per day for 10 days) in a cylindrical apparatus to accustom the rats to forage for chocolate pellets (Bio-Serv, Flemington, NJ) before the subsequent training on the circular track. Animals were screened in cohorts (2-3 young and aged rats at a time) from several independent runs of the water maze tests to total 4 Y, 7 AU, and 7 AI.

A custom-built recording hyperdrive that contained 15 independently moveable tetrodes was surgically implanted over the right hemisphere. To optimize the drive placement, recordings were performed during the surgery to find the lateral edge of CA3. The most lateral tetrode placement ranged from 3.7 to 4.2 mm lateral to bregma and from 3.4 to 4.0 posterior to bregma. The tetrode array was configured as an oval bundle, approximately 1.2 mm in length and 0.9 mm in width, with a 3 x 6 array of tetrodes spaced approximately 300 μm apart. The array was implanted at an angle of ∼35°, which is approximately parallel to the transverse axis of the dorsal hippocampus.

### Local-global cue-conflict manipulation

All rats were trained to run in a clockwise direction on a circular track for randomly placed chocolate pellets for 2-3 weeks while the tetrodes were being lowered to CA3. The circular track (76 cm outer diameter, 56 cm inner diameter) with salient local texture cues was placed in a black-curtained room with salient global cues. The cue-conflict manipulation experiment was conducted for 4 days, with 5 sessions each day. The rat was disoriented between each session. Three standard (STD) sessions (local and global cue relationships remained constant) were interleaved with two mismatch (MIS) sessions (local and global cues were rotated by equal increments but in opposite directions, producing mismatch angles of 45°, 90°, 135°, and 180°). Mismatch angles were chosen in pseudorandom order such that each angle was chosen once during the first 2 days of recording and once again during the second 2 days. On each recording day, baseline sessions (∼15 min) in which the rat rested in the holding dish were recorded prior to the start and at the end of the experiment, in order to compare recording stability before and after the experiment. Because 45° mismatch is a minor degree of separation and the cell responses in 45° MIS session do not differ markedly from STD sessions (see Lee et al., 2015), cells recorded during 45° MIS sessions were not included in this analysis.

### Two-shape experiment

During the 4 days of the cue-conflict manipulation task, subsets of the rats (3 Y, 6 AU, 6 AI) were trained in the two shapes [square (67 cm x 67 cm) and circle (76 cm diameter)] placed at a fixed location in room A. Animals were recorded in cohorts, consisting of 2-3 young and aged rats. The two-shape and two-room experiments were performed starting with the second cohort. The first cohort, which consisted of 1 Y, 1 AU, and 1 AI, did not get the two-shape and the two-room experiments. On day 1, rats were trained to forage for chocolate pellets for 20 min once in the circle enclosure and once in the square enclosure. On days 2 - 4, rats were trained for 10 min each session in the square and the circle for a total of 4 sessions in a random order (2 sessions in each shape). A striped cue card was placed on the north wall inside the chambers, and new brown paper floor was replaced between each session. The rat was allowed to rest for 5 - 10 min on the pedestal between each session. After 4 days of training, the two-shape experiment was conducted for 2 days, 4 sessions each day. During the 2 recording days, the order of the shapes was circle-box-box-circle and box-circle-circle-box, although which order was given on the first day was random.

### Two-room experiment

After 4 sessions of the different shape experiment on each recording day, the rat was placed on a pedestal and taken to room B. Two additional 10 min sessions were given in room B, once in the square enclosure and once in the circle enclosure. Two new cue cards (brown cue card on the west wall and gray hexagon shaped cue card on the east wall) were placed inside the enclosures, and a new white paper floor was replaced between each session. The order of the square and circular shape was random. There was no prior training in room B.

### Electrophysiological Recordings

Tetrodes were made from 17-μm platinum-iridium wires (California Fine Wire Co., Grover Beach, CA), with impedance reduced to ∼120 kOhms by electroplating with platinum black. Neural signals were recorded using a 64-channel wireless transmitter system (Triangle Biosystems International, Durham, NC) and transmitted to a Cheetah Data Acquisition System (Neuralynx, Bozeman, MT). The signals were amplified 1,000-5,000 times and filtered between 600 Hz and 6 kHz (for units) or 1 and 600 Hz for local field potentials (LFPs). The spike waveforms above a threshold of 40-70 μV were sampled for 1 ms at 32 kHz, and LFPs were continuously sampled at 32 kHz. The rat’s position was tracked with an overhead camera recording light emitting diodes (LEDs) position over the head of the rat (red LEDs in front and green LEDs behind) at 30 Hz.

### Data Analysis

#### Unit Isolation

Multiple waveform characteristics (e.g., spike amplitude and energy) were used to isolate single units using custom-written manual cluster-cutting software. Cells were assigned to subjective isolation quality scores from 1 (very good) to 5 (poor), depending on the cluster separation from the other clusters and from the background activity level. The isolation quality scores were made completely independently of any behavioral correlates of the cells. Using a K-means clustering analysis on three parameters (spike width, mean firing rate, and burst index), cells were classified as putative pyramidal cells (low-rate broad spikes, bursty) or putative interneurons (high-rate, narrow spikes, nonbursty) for each session (Lee et al., 2021). If a cell was classified as a putative interneuron (i.e., a high rate cell) in one of the sessions, that cell was classified as a putative interneuron for all sessions. Only well-isolated putative pyramidal cells (with isolation quality scores 1-3) were included in the analysis. Putative interneurons were excluded from the analysis.

#### Rate Maps and Place Fields

The position and the head direction of the rat were based on tracking the LEDs on the headstage connected to the hyperdrive. Analysis was performed on data restricted to times when the animal was moving forward at a speed > 3 cm/s. For 2D rate maps, the x and y dimensions of the camera image (640 x 480 pixels) were divided into bin sizes of 10 pixels (2.5 cm square bins). To create a firing rate map of a cell, the number of spikes in each bin was divided by the amount of time the rat spent in that bin. Rate maps for the circle-square environments were smoothed using the adaptive binning method (Skaggs et al., 1996); rate maps for the circular track were smoothed with a Gaussian filter (σ = 3°). For the cue-conflict manipulation data, analysis was performed on data restricted to times when the animal’s head was within the boundaries of the circular track, and the circular 2D rate maps were transformed into linear rate maps by converting the rat’s Cartesian position into units of degrees on the track. Linearized rate maps were divided into 360 bins (1°/bin). Rate maps were used to calculate the spatial information score (Skaggs et al., 1996), the mean and the peak firing rates. Place cells were identified as neurons with spatial information scores > 0.75 bit/spike, spatial information significance p < 0.01, and number of velocity-filtered spikes > 50. Spatial information significance was calculated with a shuffling procedure, in which the spike train and the position train were shifted relative to each other by a random time (minimum 30 s), the rate map was recalculated, and a spatial information score was calculated. This procedure was performed 1,000 times, and the cell was considered significant at p < 0.01 if the true information score exceeded the values of more than 990 scores from the shuffling procedure. The peak firing rate was the maximum firing rate in the rate map, and the mean firing rate was calculated by dividing the number of spikes by the duration of the session.

#### Rotational Analysis

The rotation angle of rate maps between the sessions was determined for each cell that passed the inclusion criteria in both the STD and the MIS sessions. The linearized rate map in the MIS session was circularly shifted in 1° increments and correlated with the linearized rate map in the STD session. The angle producing the maximum correlation was assigned as that cell’s rotation angle.

#### Population correlation matrices

Population correlation matrices were created by forming normalized firing rate vectors for the sample of cells at each 1° bin of the track to create 360 firing rate vectors. The firing rate of each cell in each bin is normalized to that cell’s peak firing rate. The correlation matrix contains the Pearson product moment correlations values for each of the 360 x 360 firing rate vectors. A band of high correlation along the 0° main diagonal indicates that the representation maintained coherence and did not change between the two environments.

#### Population Vector Correlation

Rate maps for the entire population of cells were arranged in an x-y-z stack, where x and y represent the two spatial dimensions and z represents the cell-identity index. The distribution of mean firing rates along the z axis represent the composite population vector for a given x-y location. Population vectors for corresponding positions were correlated across pairs of sessions. Nonoverlapping x-y locations between the two shapes were excluded from the PV correlations.

#### Statistical Analysis

All statistical tests were calculated using Matlab (Mathworks, Natick, MA) and R (R core Team, 2013).

#### Linear mixed effects models

Linear mixed effects regression models were used to estimate (separately for Y, AU, and AI animals) the total correlation difference as a function of a tetrode’s distance along the CA3 transverse axis, controlling for animal’s running speed. For each tetrode recorded along the CA3 transverse axis that had at least two or more cells recorded in that tetrode, the population correlation matrices were calculated for STD and MIS sessions (for Fig. 3) and the PV correlations were calculated between the two shapes and the two rooms (for Figs. 5 and 6). The response variable was the total correlation difference calculated for each tetrode. The key predictors were the age group of the animals and the location of the tetrode along the transverse axis. In addition to the intercept that represents the fitted value for Y rats at location 0, the fixed effects in the model include: (1) two indicator variables for the contrast of AU or AI with the reference Y group (2 df); (2) a smooth function of location (natural cube spline with 2 df) representing the baseline group’s (Y rats) dependence on location; and (3) interactions of the group and location predictors (4 df) representing the differences between the AU and AI groups and the Y group in their smooth functions of location for a total of 8 df (Hastie et al., 2009). The random effects allow animals to have random intercepts and random linear slopes along the CA3 transverse axis; the random effects account for correlation among tetrodes from the same animal so that the inferences about the group differences are valid.

A Wald test was used to test the null hypothesis that the three groups shared the same total correlation curve as a function of location (Aitchison and Silvey, 1960). The Wald test statistic was compared with a X^2^ distribution with 4 df (number of groups -1 = 2) x (number of degrees of freedom for each curve = 2). The 95% confidence intervals (CIs) were calculated for the predicted curves for each group (McCullagh and Nelder, 1989). To control for confounding by running speed, each animals’ speed was included in the fixed effects as a natural cubic spline with 3 df.

#### Logistic regression models

A generalized linear mixed effects regression model with a logistic link function was used to estimate the probability of rotating and remapping cells separately for Y, AU and AI as a function of the cell’s distance along the CA3 transverse axis. A logistic link function was used because the response variable is binary (i.e., the cell was classified as either “rotate” or “remap”). We calculated the probability that a cell was rotating. The fixed and random effects in the model were specified in the same way as in the Linear Mixed Effects models. The random effects account for correlation among cells from the same animal.

A Wald test was used to test the null hypothesis that the probability of rotating cells along the CA3 transverse axis is the same across the three groups (Aitchison and Silvey, 1960). The Wald test statistic was compared with a X^2^ distribution with 4 df. The 95% CIs were calculated for the predicted probability of rotating cells along the CA3 axis for the three groups.

#### Histological Procedures

Rats were deeply anesthetized and perfused with 4% formaldehyde. Frozen coronal sections (40 μm) were cut and stained with cresyl violet. Images of the brain slices were acquired with an IC Capture DFK 41BU02 camera (The Imaging Source, Charlotte) attached to a Motic SMZ – 168 stereoscope. All tetrode tracks were identified, and the lowest point of the track was used to determine the recording position. Distance of the CA3 tetrode was measured manually from the proximal end and scaled by the total length of CA3. The normalized position of the tetrode along the CA3 transverse axis ranged from 0 to 1, with the distal end as 1.

For tetrode assignments to proximal CA3, tetrode tips that terminated only in the pyramidal cell layer were considered. Tetrode tracks that clearly terminated above or below the pyramidal layer in the proximal CA3 region were considered to be located in the hilus and not analyzed further. While only the tetrodes that clearly terminated in the pyramidal cell layer in the proximal CA3 region were considered, any possible contamination of mossy cells will unlikely affect the interpretations of the remapping cells in the proximal region because the mossy cells and proximal CA3 cells respond similarly to the local/global cue mismatch (GoodSmith et al., 2019).

## Acknowledgements

We thank Robert McMahan, Andrew Sherwood, and Kimberly Knah for assistance in running the behavior experiments and histological procedures. We thank Audrey Branch, William Hockeimer, and Ravikrishnan Jayakumar for helpful discussion. Supported by grant P01 AG009973.

## Author Contributions

H.L., conceptualization, methodology, investigation, formal analysis, writing – original draft, writing – review & editing, and visualization; A.T., N.L., investigation and writing – review & editing; Z.W., formal analysis and writing – review & editing; S.Z., supervision and writing – review & editing; M.G., supervision, writing – review & editing, and funding; J.J.K., conceptualization, supervision, writing – original draft, writing – review & editing, and funding.

---

**Supp. S1.**
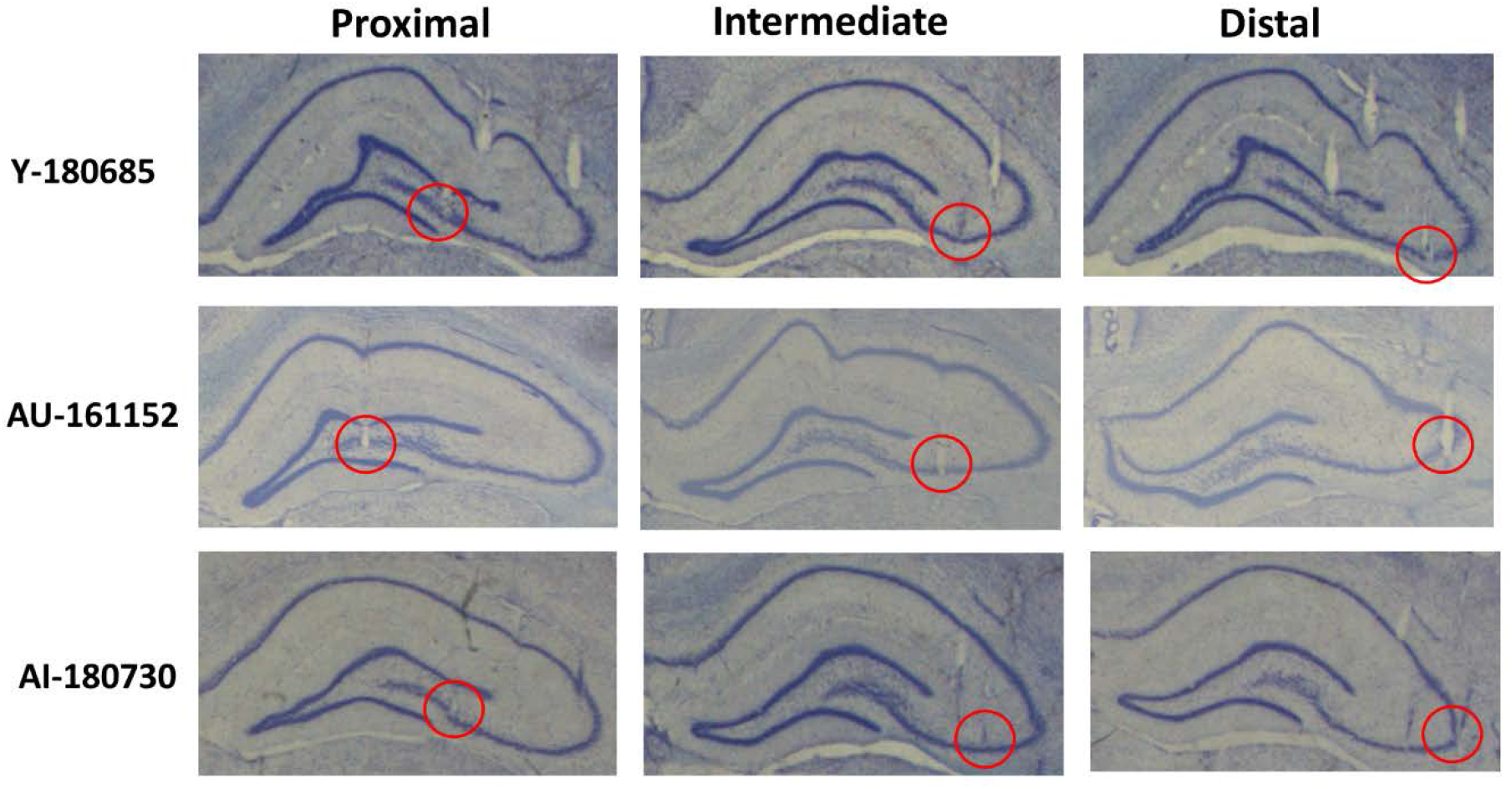
Examples of histological verification of tetrode tracks in CA3. Example rat from Y, AU, AI groups with tetrodes in all CA3 subregions. Tetrode tracks were verified on Nissl-stained coronal sections. Tetrode tips are indicated by red circles.

**Supp. S2.**
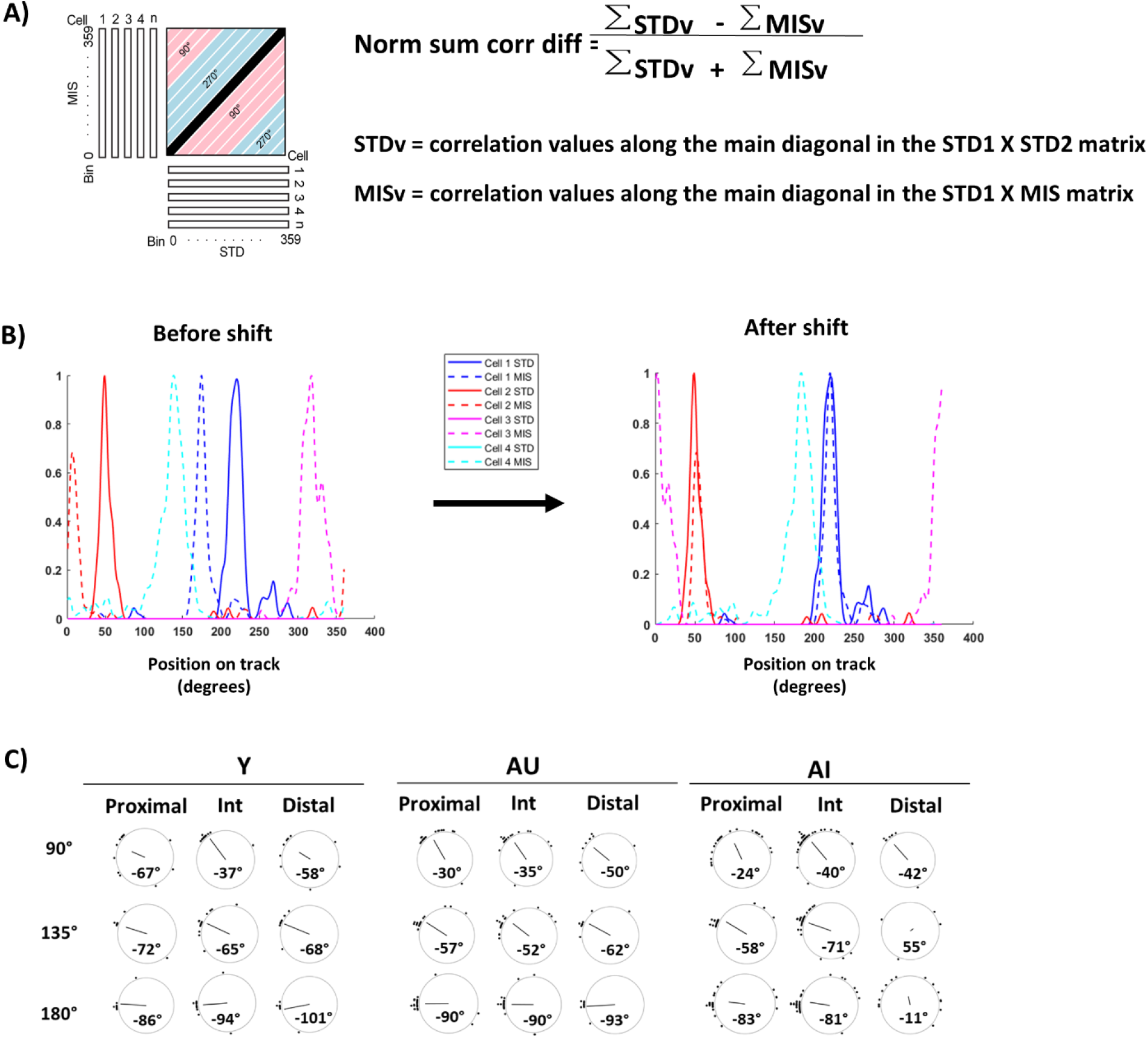
**(A)** Illustration showing the population correlation analysis for the STD vs. MIS sessions. To create these matrices, the firing rate of each cell was calculated for each 1° bin on the track and normalized to its peak rate. The firing rate maps of all *n* cells in the sample were stacked to create a 360 x *n* matrix, in which each column of the matrix represents the population firing rate vector for each angle of the track. The firing rate vectors at each angle of a STD session (STD1) were correlated with the firing rate vectors at each angle of the next STD session (STD2), to create STD1 x STD2 correlation matrices, or with the next MIS session, to create STD1 x MIS correlation matrices. The normalized correlation difference was calculated as the difference between the sum of the correlation values along the main diagonal in the STD1 x STD2 matrix (STDv) and the sum of the correlation values along the main diagonal in the STD1 x MIS matrix (MISv), divided by the sum of these two quantities. **(B)** To align the place fields in each MIS session to the place fields in the preceding STD session, each cell’s firing rate map was rotated by the mean rotation amount of each MIS session. In this example, Cell 1 (blue) and cell 2 (red) are Rotate cells (i.e., they have place fields in both STD and MIS sessions) and cell 3 (magenta) and cell 4 (cyan) are Remap cells (i.e., place fields appeared in MIS session but are silent in STD session). Before the shift (left), cell 1 fired at the 210° location on the track in the STD session (blue, solid line) but fired at the 170° location in the MIS session (blue, dotted line). Similarly, cell 2 fired at the 50° location in the STD session (red, solid line) but fired at the 10° location in the MIS session (red, dotted line). Cell 3 (magenta, dotted line) and cell 4 (cyan, dotted line) fired in the MIS session but they did not fire in the STD sessions. For this example, the mean rotation amount was 40° in the CCW direction. After shifting the place fields in the MIS session by the mean rotation amount (right), the place fields of cell 1 and cell 2 in the MIS session fired at the same location as the STD session. The place fields of cell 3 and cell 4 were also rotated by the mean rotation amount. Thus, shifting aligned the place fields of Rotate cells in the STD and MIS sessions, which allowed different experimental mismatch angles to be collapsed into one MIS session. **(C)** Mean rotation angles, indicated in bold, were close to the MIS amount in the CCW rotation, indicating a strong local cue control, except for 135 ° and 180° angles in the distal region for AI rats.

**Supp. S3.**
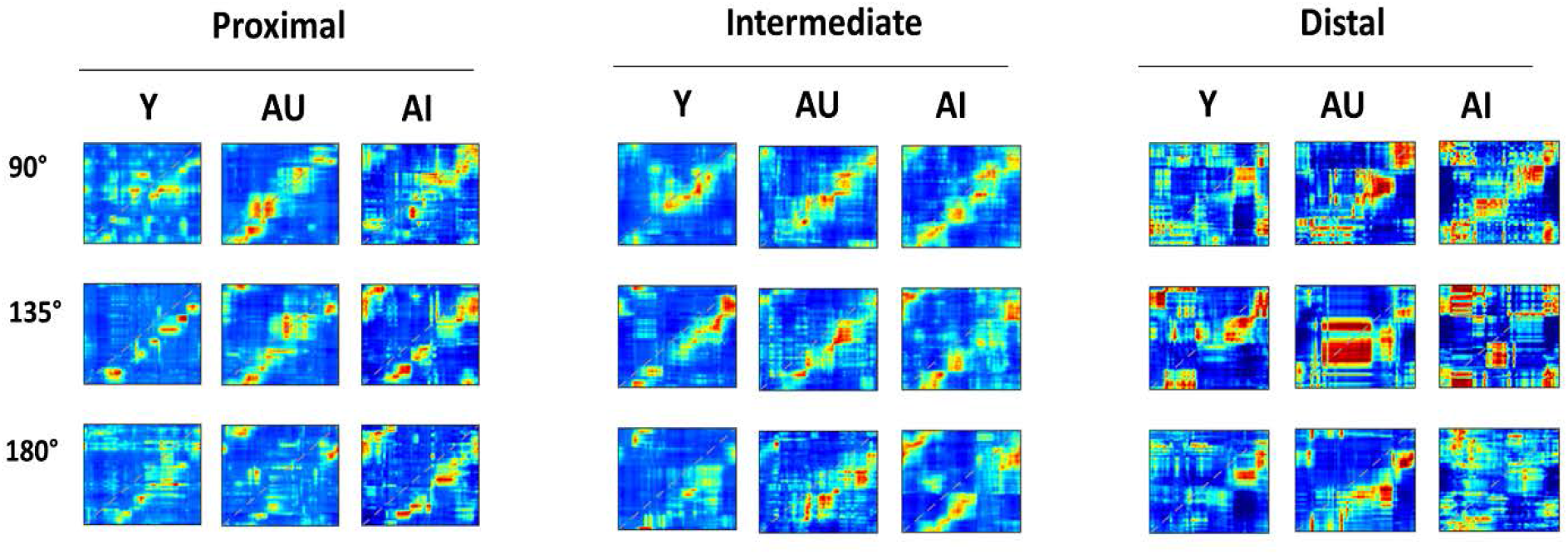
STD1 x MIS correlation matrices for each mismatch angle. In agreement with prior studies (Lee et al., 2015, 2004; Neunuebel and Knierim, 2014), the place fields tended to rotate CCW by the amount of the local-cue rotation, causing bands of higher correlation below the main diagonal (dashed line) in the STD1 x MIS correlation matrices for each mismatch amount. As the mismatch angle increased, the correlation band moved further away from the main diagonal. Note that this effect is not clearly evident in all matrices, partially due to small sample sizes from distal CA3. Because of the variability, the data from all 3 mismatch angles were collapsed for the analyses in the main text.

**Supp. S4.**
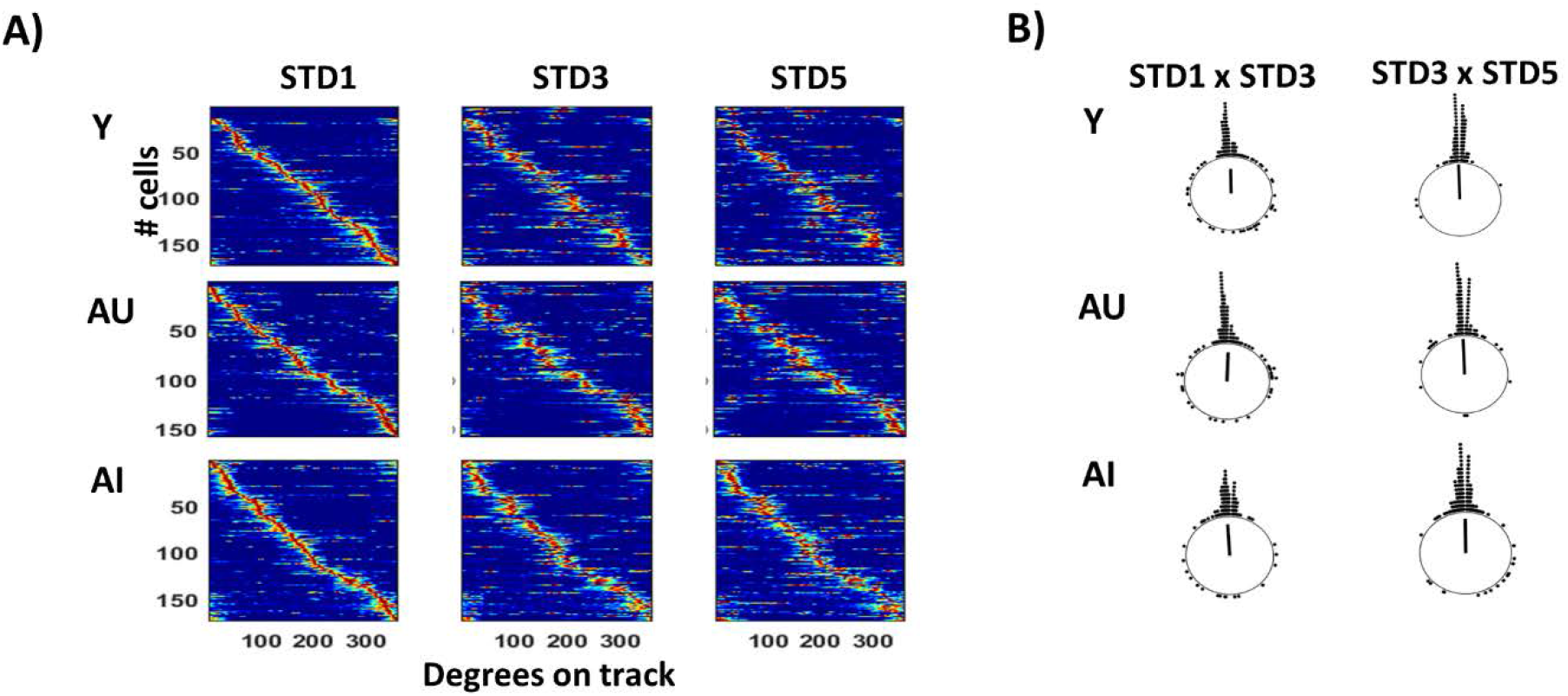
Place fields are stable between the STD sessions in all three groups. **(A)** Place cells (from all CA3 regions combined) were ordered by the peak position of their linearized rate maps in the first session (STD1) for each group. Place field ordering remained stable across the STD sessions in all groups. **(B)** To quantify stability, the angular difference in the location of each place field on the circular track between STD1 x STD3 and STD3 x STD5 were calculated. Two linearized rate maps were correlated by circularly shifting one rate map in 1° increments, and the angular difference was defined as the angle producing the maximum correlation. For all groups, the angular difference was centered around 0° with no significant group difference (Kruskal-Wallis test: STD1 x STD3: X^2^[2] = 0.43, p = 0.81; STD3 x STD5: X^2^[2] = 0.89, p = 0.64). Cells that were completely silent in one of the two sessions were excluded for the angular difference calculation (Y = 15/152; AU = 5/158; AI = 1/174).

**Supp. S5.**
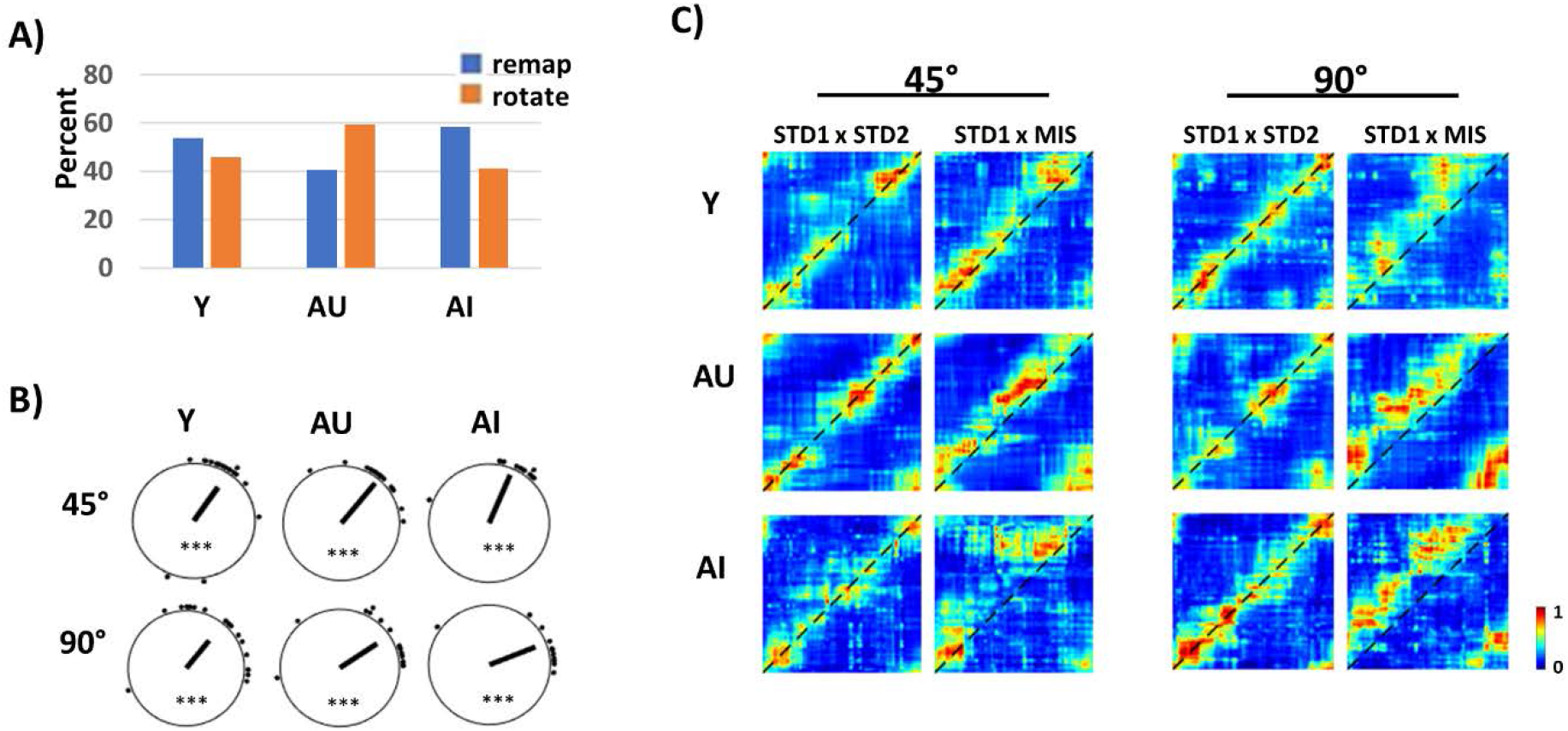
Population coherence demonstrates control by the global cues in all age groups when local cues are removed. **(A)** When local texture cues were removed and only global cues were rotated by 45° or 90°, the proportion of cells that Rotate or Remap did not differ significantly in all age groups (*X*^2^ = 4.13, p = 0.13). Cells from all CA3 subregions were pooled together. **(B)** Individual rotation amounts show significant clustering, although the rotation of the place fields was less than that of the cues, demonstrating that there were uncontrolled cues or influences of path integration that conflicted with the controlled cue set. [Rayleigh test, for 45°: Y: n = 16, z = 8.05, p < 0.0001; AU: n = 19, z = 15.76, p < 0.0001; AI: n = 12, z = 9.49, p < 0.0001; for 90°: Y: n = 19, z = 7.33, p < 0.0001; AU: n = 18, z = 10.14, p < 0.0001; AI: n = 12, z = 7.9, p < 0.0001] *** denotes p < 0.0001. **(C)** Spatial correlation matrices show that the population maintained coherence in all age groups, as indicated by the bands of high correlation. The bands of correlation in the STD1 x MIS matrices shifted upward (above the dotted diagonal line), indicating control by the global cues in all age groups.

**Supp. S6.**
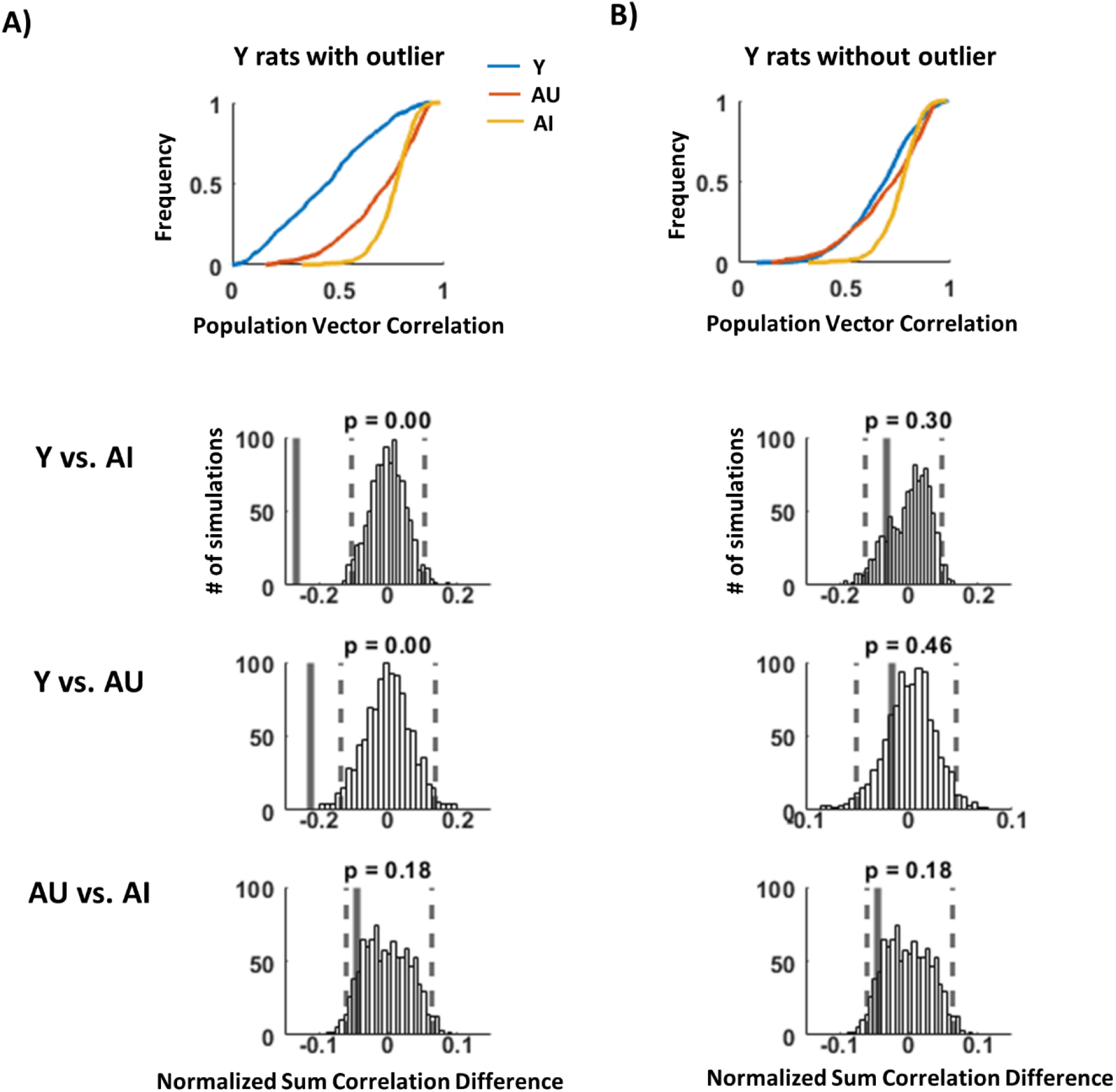
Of the 3 Y rats recorded in the two-shape experiment, 1 Y rat showed an anomalous remapping between the same shapes (i.e., the place field locations changed between the visits to the same shapes). Population vector (PV) correlations between the *same shapes* were calculated and compared between the groups with **(A)** and without **(B)** including the outlier Y rat. To measure the change in PV correlation in the same shape, the PV correlations for each of the bins (i.e., each of the values in the cumulative distribution plots) were summed for each group. To compare among the groups, the difference between the summed PV correlations of both groups was divided by the sum of the two quantities. The statistical significance was calculated by a shuffling procedure in which the cells from both groups under comparison were shuffled and randomly reassigned to each age group. The PV correlation difference was calculated from the shuffled data set, and this process was repeated 1000 times. **(A)** Cumulative distribution plots for the PV correlations between the same shape (top). When the outlier Y rat is included, PV correlations for Y rats was less correlated compared to AU and AI rats. Shuffling analysis comparing the PV correlation differences between the animal groups shows that Y rats were significantly different from AI and AU rats, while there was no difference between AU and AI rats (bottom). The observed value (thick, black line) was compared to the distribution produced by 1,000 random shuffles of the data. Dashed lines indicate the 2.5^th^ and 97.5^th^ percentiles of the distribution. **(B)** Cumulative distribution plots for the PV correlations between the same shape (top). When the outlier Y rat is excluded, PV correlations for Y rats was similar to AU and AI rats. Shuffling analysis comparing the PV correlation differences between the animal groups shows that there was no significant difference among the groups (bottom).

**Table S1.** Individual rat responses to the cue-mismatch manipulations. The CA3 analyses in the main text combined data across multiple sessions from 4 Y, 7 AU, and 7 AI rats. The total number of cell responses to the cue-mismatch manipulations observed from tetrodes in proximal, intermediate, and distal subregions for individual rats are listed.

|  | Proximal |  | Intermediate |  | Distal |  |
| --- | --- | --- | --- | --- | --- | --- |
| Rat | Remap | Rotate | Remap | Rotate | Remap | Rotate |
| Y-160281 | 6 | 3 | 3 | 3 | 6 | 14 |
| Y-160517 | 7 | 9 | 8 | 7 |  |  |
| Y-180576 | 52 | 7 | 45 | 19 |  |  |
| Y-180685 | 34 | 12 | 19 | 11 | 25 | 11 |
| AU-151055 | 9 | 13 | 9 | 16 |  |  |
| AU-160782 | 2 | 1 | 4 | 3 |  |  |
| AU-161152 | 19 | 1 | 18 | 17 | 0 | 2 |
| AU-170654 | 10 | 7 | 6 | 8 | 12 | 18 |
| AU-170711 | 14 | 11 | 3 | 2 | 1 | 5 |
| AU-190230 | 10 | 14 | 11 | 10 | 5 | 3 |
| AU-190268 | 17 | 14 | 1 | 1 |  |  |
| AI-151170 | 8 | 21 |  |  |  |  |
| AI-161201 | 2 | 5 | 5 | 6 |  |  |
| AI-170869 |  |  | 2 | 10 | 3 | 4 |
| AI-180730 | 2 | 16 | 4 | 8 | 0 | 1 |
| AI-180821 |  |  | 0 | 1 | 6 | 10 |
| AI- 190127 | 12 | 19 | 23 | 42 | 1 | 1 |
| AI-190196 |  |  | 24 | 40 | 10 | 11 |

**Table S2.** Total number of cells from individual rats during the two-shape experiment (Shape) and the two-room experiment (Room). The CA3 analyses in the main text combined data across multiple sessions from 2 Y, 6 AU, and 6 AI rats. The total number of cells recorded from tetrodes in proximal, intermediate, and distal subregions for individual rats are listed

|  | Proximal |  | Intermediate |  | Distal |  |
| --- | --- | --- | --- | --- | --- | --- |
| Rat | Shape | Room | Shape | Room | Shape | Room |
| Y-160517 | 10 | 12 | 3 | 6 |  |  |
| Y-180685 | 23 | 29 | 14 | 13 | 9 | 12 |
| AU-151055 | 17 | 16 | 15 | 19 | 2 | 2 |
| AU-161152 | 14 | 11 | 6 | 9 |  |  |
| AU-170654 | 7 | 6 | 5 | 7 | 17 | 18 |
| AU-170711 | 6 | 5 | 1 | 1 | 2 | 3 |
| AU-190230 | 6 | 9 | 15 | 16 | 2 | 4 |
| AU-190268 | 20 | 13 | 7 | 7 |  |  |
| AI-151170 | 12 | 15 |  |  |  |  |
| AI-161201 | 4 | 6 | 5 | 5 |  |  |
| AI-170869 |  |  | 9 | 4 | 1 |  |
| AI-180730 | 10 | 7 | 2 | 2 |  |  |
| AI- 190127 | 13 | 14 | 52 | 53 | 2 | 2 |
| AI-190196 |  |  | 16 | 16 | 5 | 3 |

